# GermRL: Alleviating The Germline Bias In Autoregressive Antibody Language Models Through Reinforcement Learning

**DOI:** 10.64898/2026.06.08.730660

**Authors:** Laurent Ludwig, Michael Chungyoun, Jeffrey J. Gray

## Abstract

Antibodies are powerful therapeutics whose antigen specificity arises from sequence diversity shaped during development. Recently, language models trained on large antibody repertoire datasets have enabled the generation and screening of novel candidates, but these models retain a strong germline bias. As AI adoption increases in therapeutic workflows, it is crucial to develop models that harness the diversity of antibodies necessary for the discovery of mutations that encode desirable properties. Previous work explored the germline bias in masked antibody language models, yet the bias in generative autoregressive language models has not yet been addressed. Here, we present GermRL, a lightweight and modular reinforcement learning (RL) framework capable of alleviating the germline bias in pre-trained antibody autoregressive language models through group relative policy optimization (GRPO). GermRL achieves consistent one-shot generation of antibodies that satisfy specified mutation thresholds from germline while maintaining structural plausibility. Under the lowest and highest mutation thresholds tested (5 and 35 mutations from germline), GermRL scores 0.992 and 0.950 pass@1, respectively, compared to 0.398 and 0.034 for the pre-trained language model. Within GermRL, we introduce a key pair of modifications to GRPO that increase training efficiency by discouraging reward hacking under our antibody application. Furthermore, comparison of RL generated and natural antibody sequences reveals how RL based optimization can explore alternative evolutionary mutational patterns and residue compositional strategies while preserving key global properties of natural antibodies, including identifiable germline assignments, embedding-level similarity and comparable developability profiles. Thus, RL-trained generative models optimized to promote antibody mutations through diversity from germline provide a promising framework for navigating the antibody sequence landscape, enabling exploration of novel yet biologically plausible candidates for therapeutic design.

## 1 Introduction

Antibodies are valuable therapeutics due to their antigen specificity to target relevant intervention points in disease. Antibody binding is mediated by the complementarity-retermining region (CDR) loops in the variable region of the protein, where these loops interact with the antigen through electrostatic, hydrogen-bonding, van der Waals, and hydrophobic interactions [1].

To generate antibodies capable of binding foreign antigens, naïve germline antibodies undergo V(D)J recombination, where V, D, and J gene regions are spliced together during B-cell development to form the variable region of the antibody. Antibody affinity is further refined through somatic hypermutation (SHM) after antigen exposure. Combinatorial diversity and affinity maturation adapt the immune response to protect against evolving threats.

The rapidly advancing field of artificial intelligence is transforming antibody engineering by enabling the generation and optimization of therapeutic candidates with favorable properties. Generative models have led to antibodies in a variety of formats, including VHHs, scFvs, and Fvs, that target diverse antigen epitopes ranging from infectious agents such as intestinal pathogens, influenza, RSV, and SARS-CoV-2 to endogenous targets including IL-7, TNF*α*, and GPCRs [2–17]. Central to many of these advances are protein language models (PLMs), which learn high-dimensional representations of amino acid sequences that capture structural, functional, and evolutionary relationships [18], enabling the design of proteins with improved stability or function [19–21]. In antibodies, both general PLMs and antibody-specific language models (AbLMs), which are trained on large-scale antibody sequence datasets, have been applied in zero-shot settings to guide mutational selection and improve stability and antigen-binding affinity [22, 23]. PLMs have been leveraged to train downstream predictive models for key developability properties, including clearance [24], viscosity [25], and in vitro assay performance [26–28], further establishing language-model-based approaches as a foundational technology for AI-driven antibody discovery and engineering.

To train antibody language models (AbLMs), one of the most comprehensive data sources is the Observable Antibody Space (OAS) [29], which contains more than 2 billion unpaired antibody sequences. ProGen2-OAS, trained on OAS unpaired sequences, is a representative autoregressive AbLM that generates foldable antibody sequences while respecting biological constraints such as the length of CDR loops [30].

Olsen et al. identified a germline bias in antibody language models where masked AbLMs, trained to infer masked residues from sequence context, assign higher confidence to germline residues over non-germline (NGL) residues [31]. Deposited sequences in OAS mainly derive from naive and unsorted B-cells, where the majority hold few NGL residues outside CDR3 [31]. Since language models approximate their training distribution, AbLMs inherit the germline bias present in OAS. A higher mutation load (more NGL residues) can be achieved in autoregressive language models by increasing the temperature, but this strategy often degrades the folding plausibility of the generated antibody sequence [21].

### 1.1 Related work

To reduce the germline bias in masked language models, Olsen et al. developed Ab-Lang2. Ab-Lang2 uses a focal loss to upweight NGL residues, applies alternate masking strategies, scales model depth, and pre-trains on unpaired sequences before fine-tuning on paired sequences [31].

Ab-Lang2’s framework successfully lowered NGL perplexity through data pre-processing and pre-training a model from scratch, but these strategies have not been applied to autoregressive language models. A germline bias in autoregressive models limits the possibility of sampling non-germline residues with desirable therapeutic properties.

Prior work has shown that RL can guide language models to explore beyond their pre-training distribution [32], [33], [34]. RL methods, including direct preference optimization (DPO) [35], policy proximity optimization (PPO) [36], and group relative policy optimization (GRPO) [37], have been applied to optimize PLMs for protein designability and activity, with GRPO showing strong exploration capability [38]. RL has been applied to antibodies for infilling tasks that improve biophysical characteristics like secondary structure content, solvent accessible surface area (SASA), and predicted RMSD, [39], as well as for mutation sampling to minimize changes in binding affinity [38].

### 1.2 Our contribution

We present GermRL, a customizable and modular RL framework applied to autoregressive AbLMs to alleviate the germline bias without the need for data processing or pre-training.

We train an AbLM with GermRL and evaluate its capability to generate antibodies farther from germline while maintaining high predicted structural confidence of the sequences. We introduce two key modifications to the GRPO algorithm that improve robustness to reward hacking and promote greater sequence diversity under our specific antibody application.

To assess knowledge gained during RL training, we compare in silico biophysical features of RL-generated and OAS repertoire sequences, testing whether RL preserves antibody-like properties while expanding mutational diversity beyond germline constraints.

## 2 GermRL for autoregressive antibody language model optimization

To alleviate the germline bias in autoregressive (AR) AbLMs, GermRL trains an AbLM policy by rewarding sequences above a minimum number of mutations from germline, characterized by the Levenshtein distance (LD) (**Appx. A.1**), while maintaining structural plausibility.

GermRL’s reward function combines LD and predicted local distance difference test (pLDDT from ESMfold [20]) to reward sequences that exceed both a set LD and pLDDT threshold (LD_thresh_ and P_thresh_) as well as satisfy the following conditions: length greater than 100 residues, identifiable human V and J germline genes, and only canonical amino acid tokens. The complete reward function is described in **Appx. A.2**.

Our GRPO implementation follows the on-policy and outcome-supervised RL scheme of Shao et al.[37] (**Appx. A.3**). Building on prior work, we incorporate two additions before applying our own modifications: (1) an entropy loss term to promote residue diversity [40] and (2) the “clip higher” strategy from dynamic sampling policy optimization (DAPO) for asymmetric clipping [41] (**Appx. A.4**). We refer to this configuration as “default GRPO”.

### 2.1 Applying antibody sequence prompts to GRPO

To promote exploration of diverse sequences at high LD, we generate sequences without constraining our AbLM to specific starting 3-residue prompt anchors. Unlike standard GRPO, which samples prompts from a dataset of queries [37], we frame the task as unconstrained sequence generation by initializing all generations from a single <start> token.

## 3 Default GRPO induces entropy collapse at higher LD thresholds

### 3.1 Training setup

Using the ProGen2-OAS implementation on Hugging Face by Hrbáň et al. [42], adapted from Nijkamp et al. [30], we create a series of four ProGen2-RL policies (LD5+, LD15+, LD25+, and LD35+) at increasing LD_thresh_ set in the GermRL reward function (**Appx. A.2**). For example, LD5+ denotes ProGen2-OAS trained with a LD threshold of 5, such that sequences exceeding 5 mutations from germline receive the maximum reward. At all LD thresholds, the pLDDT needed to reach the maximum reward is 0.7, a number that represents a generally confident prediction [43].

### 3.2 RL trained LM generates antibodies above LD threshold

We evaluate whether ProGen2-RL policies can generate sequences far from germline while retaining foldability by comparing the LD and pLDDT of 1,000 generated antibodies from each RL policy and base ProGen2-OAS at multiple temperatures (**Fig. 1a, Fig. 1b**).

**Figure 1:**
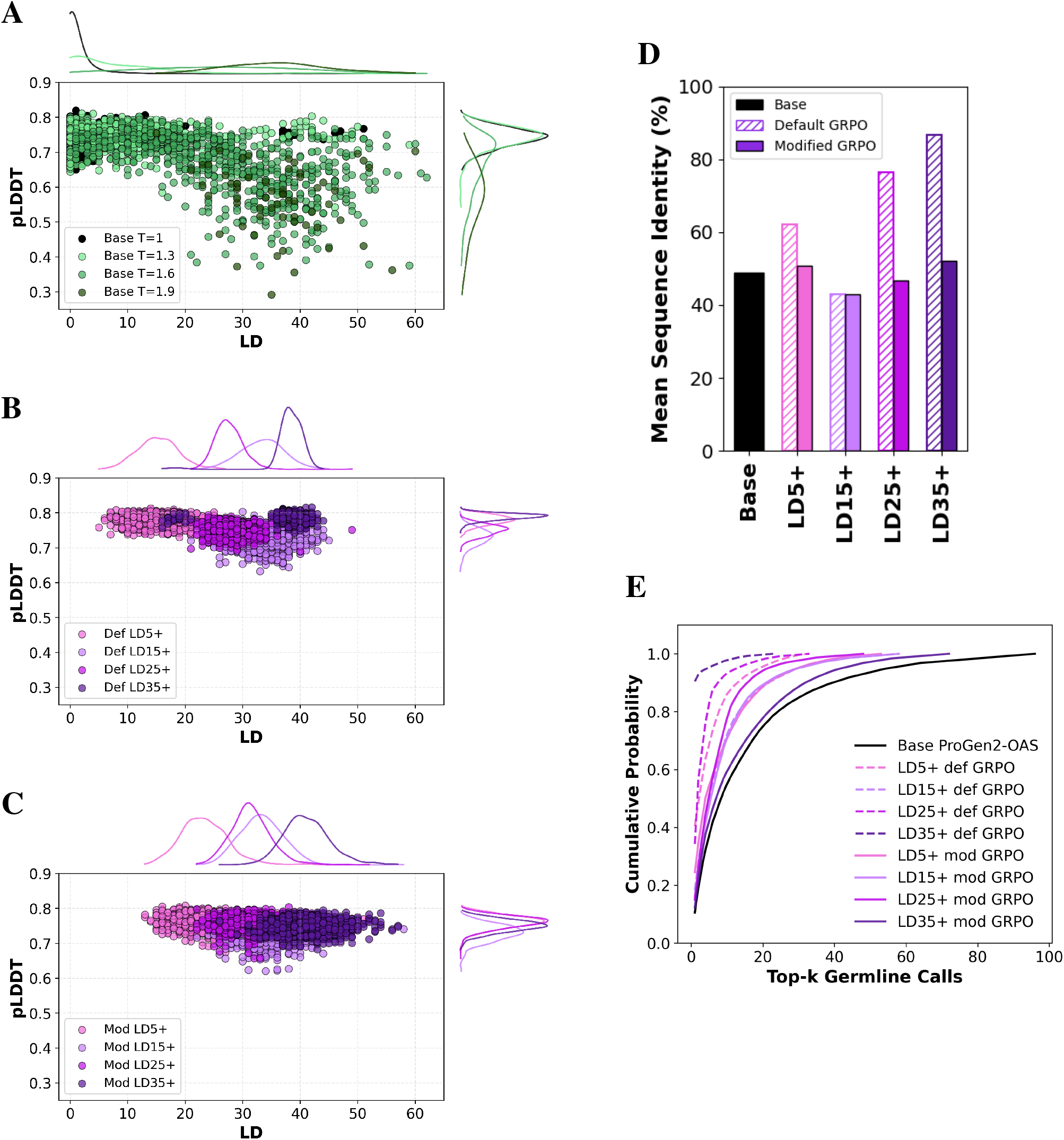
**(A)** LD and pLDDT of base ProGen2-OAS generations for each temperature. **(B)** LD and pLDDT of ProGen2-RL policy (trained with default GRPO) generations for each LD threshold set in GermRL. **(C)** LD and pLDDT of ProGen2-RL policy (trained with modified GRPO) generations for each LD threshold set in GermRL. **(D)** Mean sequence similarity between each pair of antibodies from all base ProGen2-OAS (T = 1.0) generations and RL policy generations from training with default and modified GRPO. **(E)** Cumulative distribution of top-*k* germline V calls (including alleles) of generated sequences from base ProGen2-OAS (T = 1.0) and after RL training with default and modified GRPO.

Base ProGen2-OAS at the default sampling temperature (T = 1.0) generates sequences clustered near zero LD. Increasing the sampling temperature shifts the generated sequences toward higher LD values while reducing pLDDT (**Fig. 1a**). In contrast, ProGen2-RL policies, trained with GRPO, generate antibodies with substantially higher LD while maintaining pLDDT values above the 0.7 threshold. RL policies trained at higher LD thresholds generate sequences further from germline (**Fig. 1b**). Furthermore, in base ProGen2-OAS (no RL), elevated sampling temperatures increase the proportion of sequences with no identifiable germline assignment. Specifically, 5% of sequences lack an identifiable germline at T = 1.0, increasing to 16% at T = 1.3, 37% at T = 1.6, and 94% at T = 1.9 (**Appx. B**). In contrast, all RL-trained policies produced by GermRL generate antibodies with identifiable germline assignments across the evaluated conditions (**Appx. B**).

These results indicate that base ProGen2-OAS exhibits a germline bias. Greater variability in token sampling, induced by temperature, increases diversity from germline but diminishes the structural plausibility and antibody-like characteristics learned by the LM. Our RL framework successfully generates antibodies at high LD while maintaining a foldable and identifiable antibody sequencee. Furthermore, incorporating a tunable LD_thresh_ in the GermRL reward function explicitly guides the policy toward generating sequences that exceed the desired LD threshold.

To evaluate the consistency of ProGen2-RL policies in generating proper antibodies far from germline, we use the pass@1 metric, defined as the probability that a single generation attempt yields a successful sequence (**Appx. C**). We first define success as generating sequences with LD *>* LD_thresh_ (top four rows, **Table 1**), while satisfying the length, human germline, and canonical amino acid token constraints. We then apply a stricter success criterion requiring sequences to satisfy both LD_thresh_ and P_thresh_ (bottom four rows, **Table 1**).

**Table 1.** Probability of a single generation success (pass@1) is shown for the LD only success condition in the top four rows and LD with foldability success condition in the bottom four rows across default and modified GRPO trained policies with GermRL and base ProGen2-OAS at different temperatures. Bolded entries represent the highest pass@1 value for each LD threshold between the corresponding RL trained policy and base ProGen2-OAS at varying temperatures.

| Condition | LD<sub>thresh</sub> | GermRL (Default GRPO) |  |  |  | GermRL (Modified GRPO) |  |  |  | Base ProGen2-OAS Temp |  |  |  |
| --- | --- | --- | --- | --- | --- | --- | --- | --- | --- | --- | --- | --- | --- |
|  |  | LD5+ | LD15+ | LD25+ | LD35+ | LD5+ | LD15+ | LD25+ | LD35+ | 1.0 | 1.3 | 1.6 | 1.9 |
| LD > LD<sub>thresh</sub> | 5 | **0.999** | 0.991 | 0.999 | 1.000 | 0.996 | 0.990 | 0.995 | 0.994 | 0.068 | 0.431 | 0.596 | 0.060 |
|  | 15 | 0.580 | **0.991** | 0.999 | 1.000 | 0.992 | 0.990 | 0.995 | 0.994 | 0.014 | 0.147 | 0.503 | 0.060 |
|  | 25 | 0.015 | 0.980 | 0.925 | 0.979 | 0.356 | 0.985 | **0.982** | 0.994 | 0.011 | 0.054 | 0.349 | 0.054 |
|  | 35 | 0.000 | 0.398 | 0.005 | **0.979** | 0.013 | 0.359 | 0.162 | 0.960 | 0.011 | 0.036 | 0.182 | 0.035 |
| LD > LD<sub>thresh</sub><br>and pLDDT > 0.7 | 5 | **0.999** | 0.878 | 0.993 | 1.000 | 0.992 | 0.866 | 0.984 | 0.984 | 0.067 | 0.398 | 0.232 | 0.003 |
|  | 15 | 0.580 | **0.878** | 0.993 | 1.000 | 0.988 | 0.866 | 0.984 | 0.984 | 0.014 | 0.135 | 0.158 | 0.003 |
|  | 25 | 0.015 | 0.867 | 0.920 | 0.979 | 0.354 | 0.861 | **0.971** | 0.984 | 0.011 | 0.049 | 0.069 | 0.002 |
|  | 35 | 0.000 | 0.355 | 0.004 | **0.979** | 0.013 | 0.308 | 0.162 | 0.950 | 0.011 | 0.034 | 0.027 | 0.001 |

Under both criteria, applying GermRL with default GRPO to ProGen2-OAS achieves pass@1 values close to 1 at the LD thresholds used during training, outperforming base ProGen2-OAS, whose highest pass@1 score is 0.398 under the easiest LD threshold (LD5) (**Table 1**). Pass@1 results demonstrate that ProGen2-RL policies can reliably generate high-LD antibodies meeting the desired design constraints in a single sampling attempt, substantially improving one-shot generation performance.

GermRL (Default GRPO) GermRL (Modified GRPO) Base ProGen2-OAS Temp RL is known to be vulnerable to reward hacking, where policies exploit the reward function to obtain high rewards without learning the objective [44]. Therefore, we next checked whether the successful high-LD generations arise from the policy repeatedly generating similar sequences to exploit the reward. To measure antibody generations diversity, we calculate the mean pairwise sequence identity among the 1,000 antibodies generated from each ProGen2-RL policy, and we compare the values to base ProGen2-OAS at T = 1.0 (**Fig. 1d**).

At higher LD_thresh_ values, RL-generated sequences exhibit increased sequence similarity (**Fig. 1d**). Specifically, LD25+ and LD35+ generations hold 77% and 87% sequence identity while base ProGen2-OAS generated sequences remain diverse at 49%. Sequence logos (**Fig. 2b** and **Appx. D**) reveal entropy collapse, where the policy converges to a narrow subset of sequences that maximize reward, a sign of reward exploitation under default GRPO.

**Figure 2:**
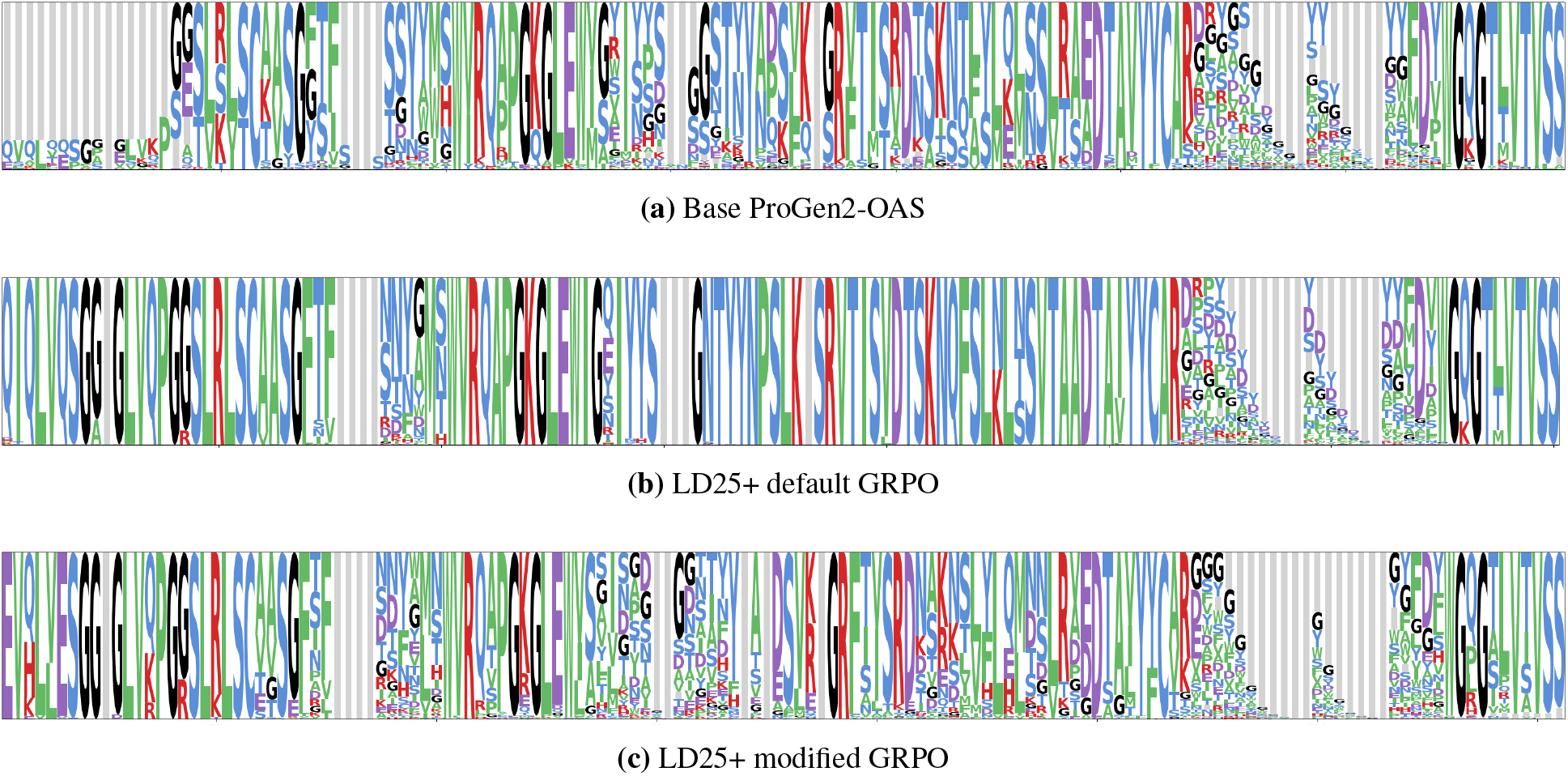
Example sequence logoplots of generations from base ProGen2-OAS and ProGen2-RL trained with default and modified GRPO at the LD25 threshold. Complete logos are shown in **Appx. D**.

To explore the entropy collapse, we analyze human germline V-gene calls (including alleles) across the 1,000 generated sequences, rank them from most to least frequent, and plot the cumulative distribution of the top-*k* germline calls (**Fig. 1e**). At high LD_thresh_, ProGen2-RL cumulative probability curves are steep, which indicates a select handful of germline V calls make up the majority of generations. Thus, RL policies exploit similar germlines, leading to the high sequence similarity of RL-generated antibodies. **Fig. 3** and **Appx. E** include piecharts to further demonstrate the divergence in germline diversity by breaking down the distribution of germline V and J calls from each RL policy compared to base ProGen2-OAS.

**Figure 3:**
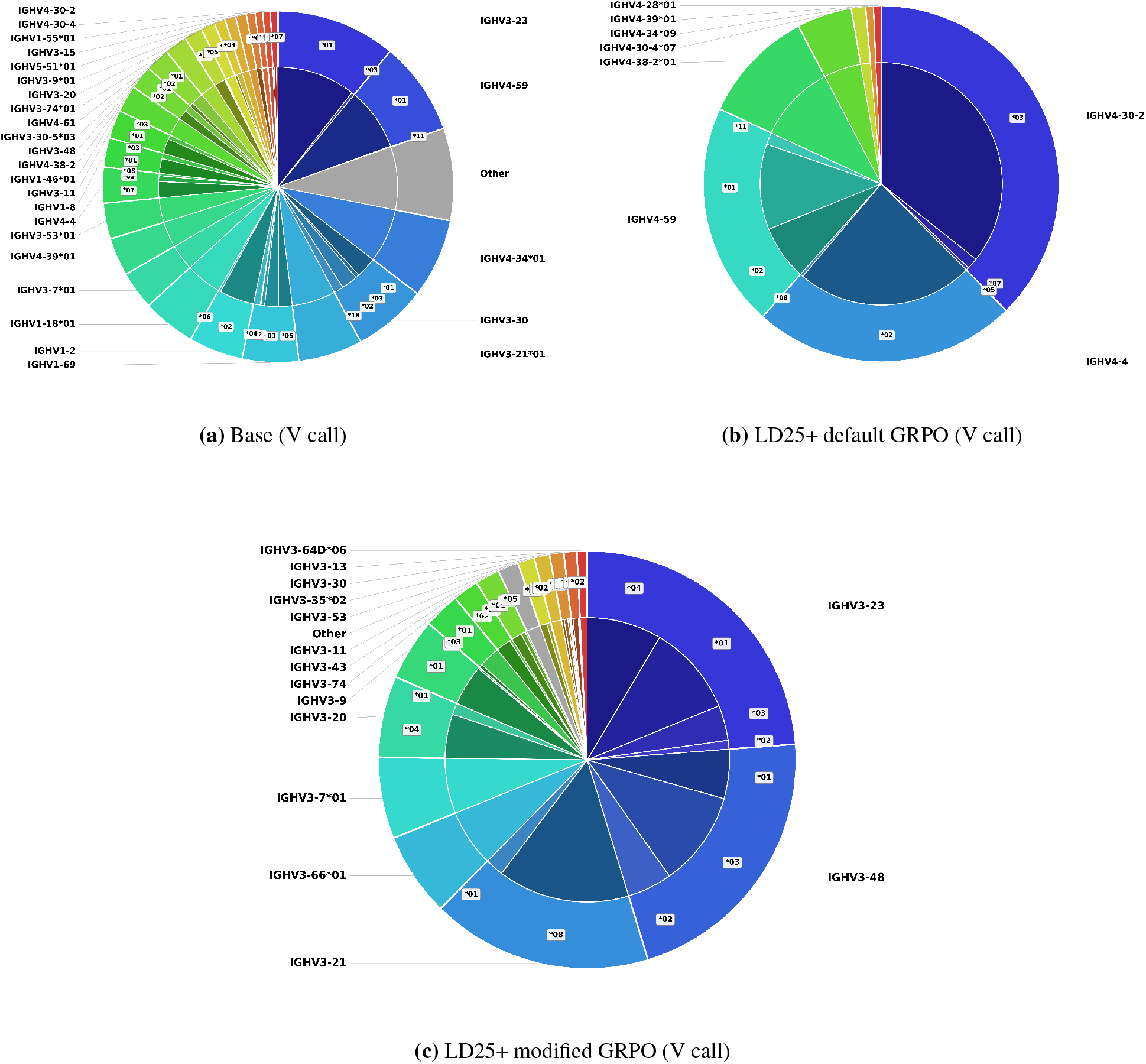
Example comparison of germline V gene usage across base ProGen2-OAS and ProGen2-RL under default and modified GRPO at the LD25 threshold. Complete pieplots for V and J calls are shown in **Appx. E**.

In GRPO for natural language tasks, the policy is exposed to diverse initial states through prompts sampled from a dataset. Because optimal completions are prompt-dependent, the policy must generalize across a range of trajectories, reducing the risk of overfitting to any single solution.

Here, we attribute the entropy collapse to the initialization of every episode from the <start> token in our training procedure. While this approach eliminates reliance on residue anchors to enable unconstrained antibody generation, it also eliminates variability in the initial state distribution. Consequently, once the policy discovers a trajectory that satisfies a high LD objective, identical starting conditions allow that trajectory to be reproduced consistently. At challenging LD objectives, the resulting reduction in exploration leads to the observed entropy collapse. Thus, we next sought strategies to modify GRPO to promote generated sequence diversity.

## 4 GRPO modifications alleviate the germline bias and reward hacking

We make two changes to address the limitations of the default GRPO algorithm:

1. **Delay updates of the old policy:**to prevent early successful actions from prematurely defining the update trust-region and driving reliance on specific trajectories, we synchronize the old policy weights with the updating policy only after all steps within an epoch, rather than after each step.
2. **Sample from the current policy, not the old:** to ensure trajectories reflect the model’s most recent improvements and preserve on-policy reinforcement learning with the first modification, we generate trajectories during each step from the updating policy rather than the old policy.

We explain both modifications in detail in **Appx. A.5**.

### 4.1 Modified GRPO achieves similar pass@1 to default GRPO while preventing entropy collapse

We train ProGen2-RL policies using modified GRPO at increasing LD thresholds (LD5+, LD15+, LD25+, and LD35+) and generate 1,000 antibodies from each policy following RL training.

To assess whether our GRPO modifications mitigate the entropy collapse, we analyze the mean sequence identity and the cumulative density of top-*k* germline calls among generated antibodies (**Fig. 1d, Fig. 1e**). Logos of the sampled residues per position are shown in **Fig. 2** and **Appx. D**, while pieplots of germline call distribution are shown in **Fig. 3** and **Appx. E**. Compared to default GRPO, modified GRPO policies reduce sequence identity, comparable with base ProGen2-OAS at T = 1.0. LD25+ and LD35+ RL policies with modified GRPO achieve 47% and 52% sequence identity, respectively. While modified GRPO cumulative probability curves are shallower than default GRPO, they remain steep relative to base ProGen2-OAS. Thus the GRPO modifications discourage reward hacking by promoting sequence and germline call diversity in generated antibodies over default GRPO. However, RL training still constrains the breadth of germline space explored compared to base ProGen2-OAS.

We next confirm whether the GRPO modifications preserve the ability to generate antibodies far from germline while maintaining folding quality by analyzing the LD and pLDDT distributions of generated antibodies (**Fig. 1c**). Modified GRPO generates antibodies with elevated LD while maintaining pLDDT distributions above the 0.7 threshold, indicating that the modified framework continues to alleviate germline bias without compromising predicted foldability (**Fig. 1c**). Furthermore, the pass@1 data indicate the modified framework maintains consistent one-shot generation performance for foldable, high-LD antibodies (**Table 1**).

## 5 RL-trained ProGen2-OAS learns to fill antibody prefix gaps through prefix grafting

AbLMs also inherit a secondary bias in generating missing FR1 antibodies that arises from training on truncated sequences due to experimental limitations in primer design [30]. A common workaround is to anchor generation with a starting 3-residue prompt such as “EVQ” [21], [30].

We initialize generations from the <start> token to enable open-ended generation without anchor conditioning, which frequently produces antibody sequences with truncated prefixes. To address this, we introduce a “prefix grafting” technique in GermRL. During training, we allow ProGen2-OAS to generate both FR1 incomplete and complete antibody sequences. We determine whether the sequence lacks a prefix by aligning it to its closest germline V call. If a prefix gap is detected, we fill the missing residues with the corresponding germline sequence and use the grafted antibody within the probability ratio for GRPO computation (**Appx. A.3**).

**Fig. 4** shows the mean fraction of trajectories requiring prefix grafting per training epoch; it converges toward zero as training progresses. Although the conditional presence of a prefix is not factored into the reward function, the policy learns over the course of RL training to generate complete antibodies, effectively eliminating this additional OAS-inherited bias.

**Figure 4:**
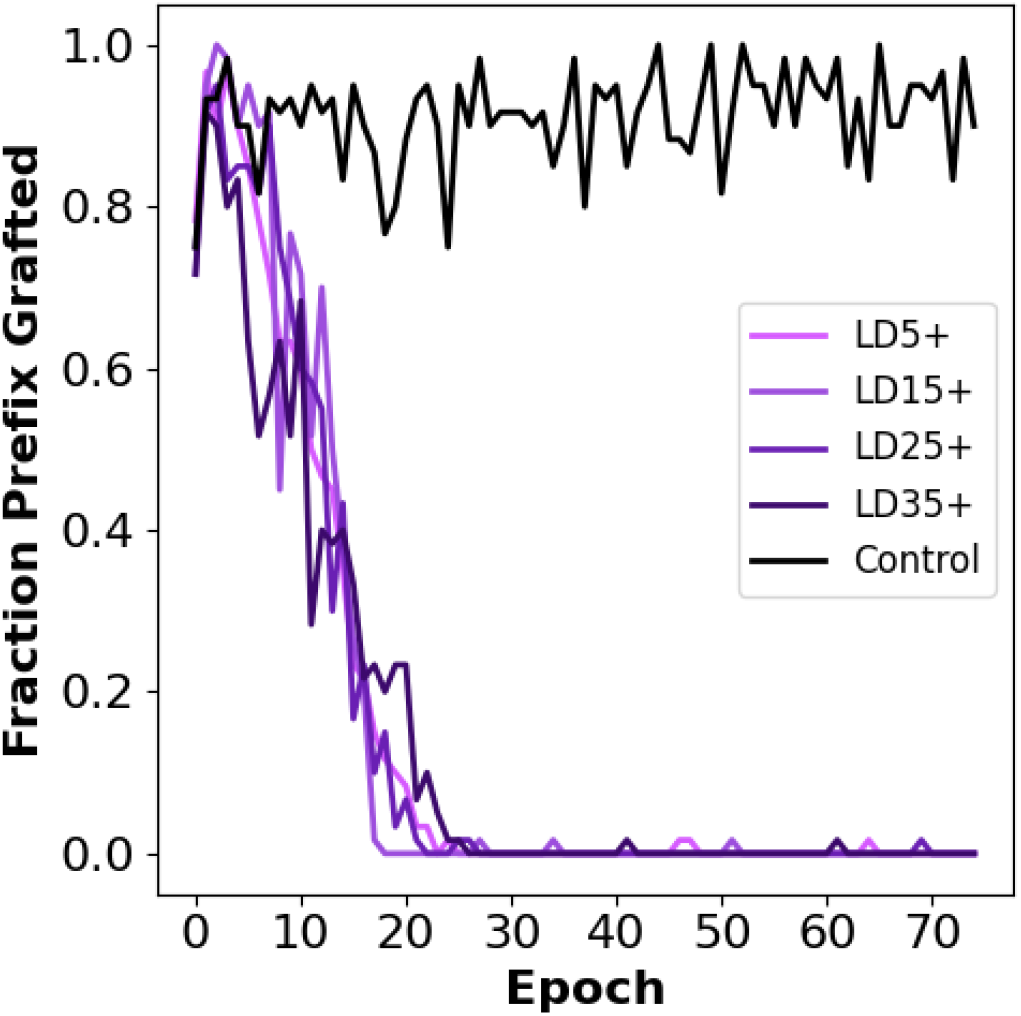
Mean fraction of generated trajectories requiring prefix grafting to fill incomplete prefixes within the training steps of each epoch during RL training with modified GRPO. Control policy represents LD25+ trained with modified GRPO without prefix grafting.

## 6 ProGen2-RL policies adopt alternate mutational strategies while preserving key biophysical antibody properties

### 6.1 Analysis framework

We next ask whether RL-trained policies learn to generate high-LD antibodies by reproducing OAS sequences or by exploring novel antibody sequence space. To assess the extent of knowledge acquired during RL training, we compare antibody properties between natural OAS and RL-generated antibodies across LD bins.

To generate antibodies within a controlled LD range, we introduce a bounded GermRL objective (**Appx. F**) to train three ProGen2-RL policies (LD5-15, LD15-25, and LD25-35). After training, each policy generates antibodies within its respective LD range (**Appx. F**).

For each LD bin, we compare 1,000 RL-generated antibodies against 5,000 natural OAS antibodies. To control for LD as a confounding variable, we match the LD distributions of sampled OAS antibodies to those of the RL-generated sequences within each bin. Because many OAS antibodies contain truncated FR1 regions, the LD may reflect different overall mutation burdens in truncated OAS sequences compared to full-length RL-generated antibodies. To account for this, we first sample complete OAS antibodies to establish the relationship between full-sequence LD and LD computed after removing FR1. We then derive 95% confidence intervals mapping truncated LD values to their corresponding full-sequence LD ranges, enabling the sampling of truncated OAS antibodies based on estimated true full-length LD to match the RL-generated distribution.

Since base ProGen2-OAS and ProGen2-RL preferentially generate heavy chain sequences (**Appx. E**), we restrict our analyses to heavy-chain OAS antibodies. Finally, to distinguish from antibody property divergence incurred through pre-training alone, we compare OAS sequences to antibodies generated from base ProGen2-OAS at T = .3 (**Appx. H**). For all OAS antibody sampling, we use seqkit [45] to randomly shuffle human antibody sequences from unpaired OAS clustered at 80% identity.

### 6.2 RL generated antibodies exhibit altered amino acid composition and mutation positioning relative to OAS antibodies

We first assess whether RL-generated antibodies exhibit divergent mutational compositions across antibody regions. We compute the fraction of residues differing from the germline sequence in FR1, FR2, FR3, FR4, CDR1, and CDR2 for both RL-generated and OAS antibodies across each LD bin (**Fig. 5**). All RL-generated antibodies contain complete FR1, while only 2% of sampled OAS sequences do. We omit CDR3 due to its intrinsic variability, which requires omission of D-gene alignment in the LD computation used for the reward function (**Appx. A.1**). We define antibody regions according to the IMGT numbering scheme [46].

**Figure 5:**
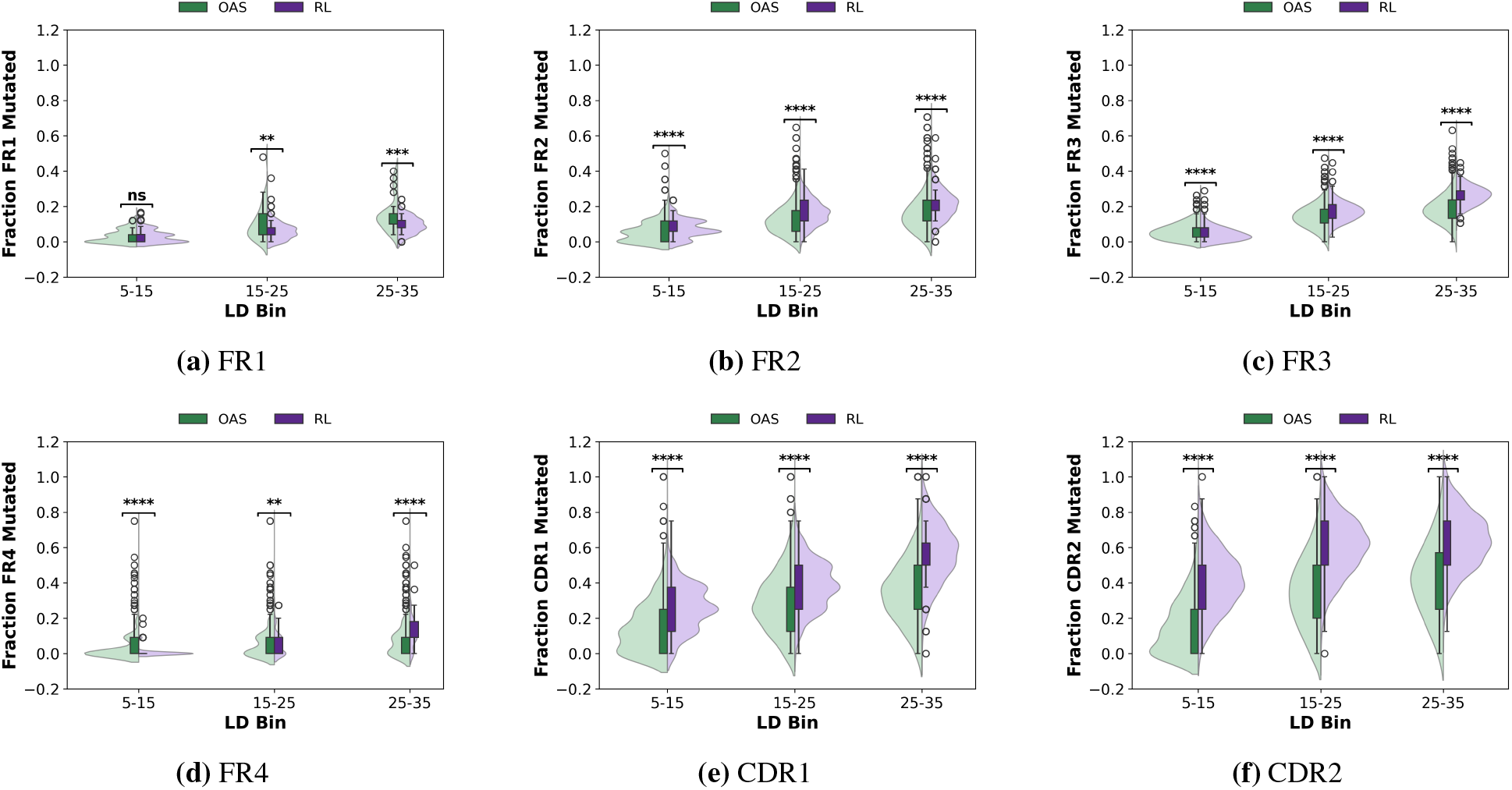
Fraction of each antibody region that is mutated for RL generated and OAS sequences across different LD bins.

Across all LD bins, RL-generated antibodies exhibit a higher fraction of mutated residues within CDR regions relative to OAS antibodies. At higher LD bins, ProGen2-RL antibodies additionally display elevated mutation frequencies in FR2 and FR3 (for both LD 15-25 and LD 25-35 ProGen2-RL policies) and in FR4 (LD 25-35 ProGen2-RL policy only). In contrast, RL-generated antibodies show reduced mutation frequencies in FR1 at high LD bins and in FR4 at low LD. (**Fig. 5**).

In comparison, mutational divergence between base ProGen2-OAS and OAS antibodies is largely restricted to CDR regions and FR3. Although base ProGen2-OAS exhibits CDR1 divergence comparable to ProGen2-RL at low LD bins, ProGen2-RL substantially amplifies CDR2 mutations across all LD bins relative to OAS (**Appx. H.1**).

These findings indicate that the mutational distribution observed in RL-generated antibodies does not arise solely from pre-training. Instead, RL training promotes a region-specific mutational strategy enriched in CDRs. Given the central role of CDRs in antigen binding, this enrichment may facilitate exploration of functionally beneficial mutations at high LD. Notably, mutational trends vary across LD bins, suggesting that the policy adopts distinct optimization strategies depending on LD difficulty.

We attribute the preferential accumulation of mutations within CDRs to the prefix-grafting approach that could bias the policy toward preserving germline-like FR1 regions to fill truncated prefixes.

We next investigate whether amino acid composition across antibody regions remains consistent between RL-generated and natural OAS antibodies. To quantify distributional differences, we compute both the absolute difference in mean fractional amino acid composition and the Kullback-Leibler (KL) divergence between RL and OAS residue frequency distributions (**Fig. 6b**).

**Figure 6:**
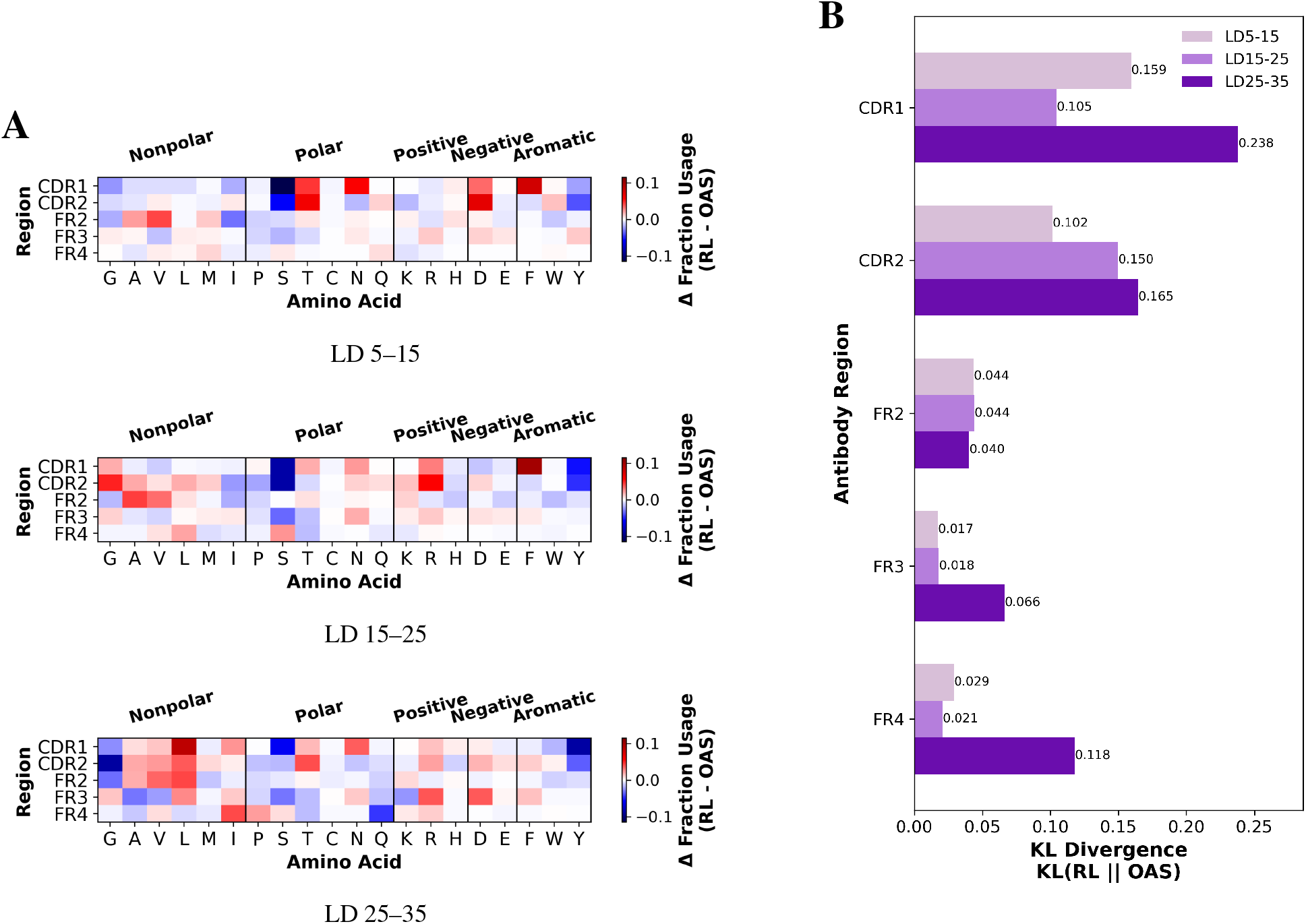
(**a**) Breakdown of difference in regional amino acid fractional composition between RL generated and OAS sequences across LD bins. (**b**) KL divergence between RL and OAS amino acid distributions for increasing LD ranges.

CDR1 and CDR2 exhibit the largest KL divergence in amino acid composition across all LD bins (**Fig. 6a**). FR2 and FR3 show relatively small divergence, while FR4 demonstrates pronounced deviation at LD25-35. KL divergence between base ProGen2-OAS and OAS sequences are significantly smaller at around a tenth of the RL and OAS divergence across all regions (**Appx. H.2c**).

At the residue level, the absolute difference in fractional composition between RL and OAS remains small, at approximately 10% or less, across all regions and LD bins (**Fig. 6a**). However, this similarity is lower than that observed between base ProGen2-OAS and OAS, for which the difference in residue fractional composition is substantially smaller across all regions (**Appx. H.2**).

Aromatic amino acids are largely conserved between RL and OAS in FR4. In CDR1, RL-generated sequences prefer phenylalanine, while OAS sequences frequently use tryptophan in both CDR1 and CDR2 over RL. For polar residues, RL-generated antibodies show consistently higher usage of threonine and asparagine in CDR1 across all LD bins, while in the same region, OAS sequences prefer serine.

Nonpolar residue composition varies across LD bins, with shifts in preference between RL and OAS, particularly for glycine in CDR1, CDR2, and FR2. RL-generated antibodies show increased usage of negatively charged residues, especially aspartate in CDR2 and FR3 across all LD bins. At LD25-35, RL-generated sequences hold an increased proportion of leucine across most regions relative to OAS.

Base ProGen2-OAS antibodies exhibit residue preference trends similar to those observed in RL-generated antibodies relative to natural OAS sequences. In particular, CDR regions show enrichment of serine, as well as increased usage of aspartic acid and asparagine, compared to OAS (**Appx. H.2**).

These findings suggest that RL regional residue preferences deviate from natural OAS sequences to a greater extent than the deviation introduced through pre-training alone. We primarily observe these differences in CDR regions and in specific residue classes, reflecting learned sequence design strategies acquired during RL training. Compositional differences vary across LD bins, showing that the RL mutation strategy evolves as the policy explores higher LD ranges. The increased KL divergence at LD25-35 across multiple regions reveals that the policy increasingly departs from natural amino acid usage patterns as it optimizes toward higher LD.

### 6.3 RL-generated antibodies exhibit lowered conservation of evolutionary substitutions

To assess whether RL-generated sequences follow natural evolutionary substitution patterns, we compute the average BLOSUM62 matrix score from mutations from germline for OAS and RL antibodies (**Fig. 7**). BLOSUM matrices score amino acid substitutions according to their evolutionary likelihood. Prior work demonstrated a correlation between BLOSUM scores and AbLM likelihood [47].

**Figure 7:**
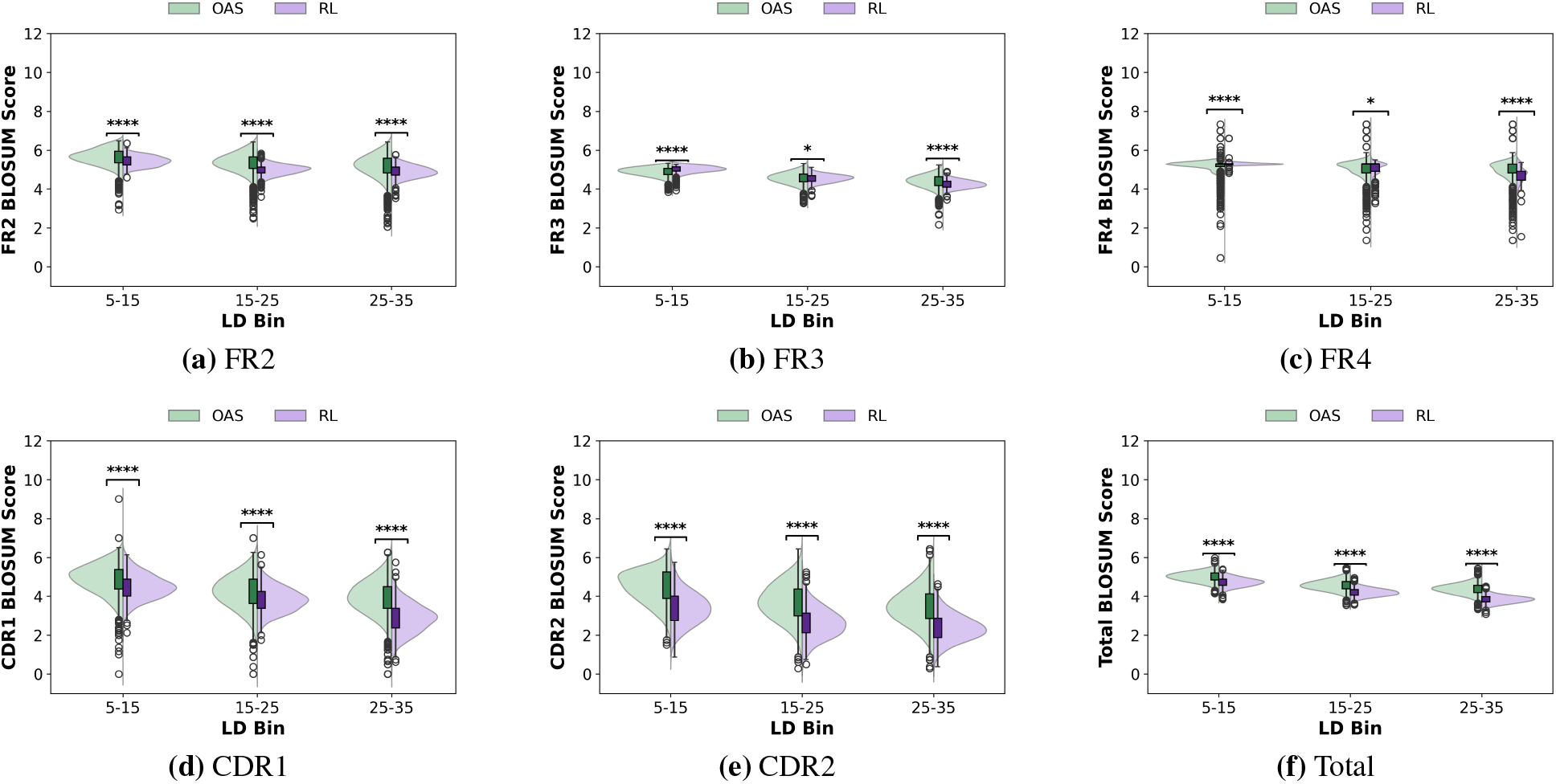
Mean BLOSUM62 scores from RL and OAS sequences across LD bins and antibody regions.

For both OAS and RL-generated antibodies, total BLOSUM scores decrease with increasing LD. Across all LD bins, ProGen2-RL antibodies exhibit lower total BLOSUM scores than OAS antibodies, with the disparity increasing at higher LD. Region-wise analysis shows lower BLOSUM scores for RL-generated sequences across most antibody regions (**Fig. 7**). Base ProGen2-OAS antibodies also exhibit reduced BLOSUM scores relative to OAS, although to a lesser extent than RL-generated antibodies (**Appx. H.3**). However, at LD 15-25, both RL and base antibodies show comparable scores to OAS in FR3 and FR4, while at LD 5-15, RL antibodies achieve higher BLOSUM scores than OAS antibodies in FR3 (**Fig. 7**).

These findings indicate that BLOSUM scores diverge at higher LD and within CDR regions of ProGen2-RL antibodies. Overall, RL-generated antibodies do not recapitulate the evolutionary substitution patterns observed in natural antibodies to a greater extent than the pre-trained AbLM.

### 6.4 ProGen2-RL preserves high-level antibody features through embedding-space semantic similarity

Given the divergent regional amino acid compositions and mutation patterns observed in RL-generated sequences, we next ask whether they still retain high-level semantic similarity to natural OAS antibodies by examining the distribution of sequence embeddings. We generate AntiBERTy embeddings [48] for both RL-generated and OAS sequences across LD bins. For each sequence, we pool the per-residue embeddings (dimension *N*× 512) to obtain a single vector of dimension 512, then apply UMAP for dimensionality reduction (**Fig. 8**). For OAS sequences with truncated FR1, we apply prefix grafting from germline to ensure embeddings reflect complete antibodies.

**Figure 8:**
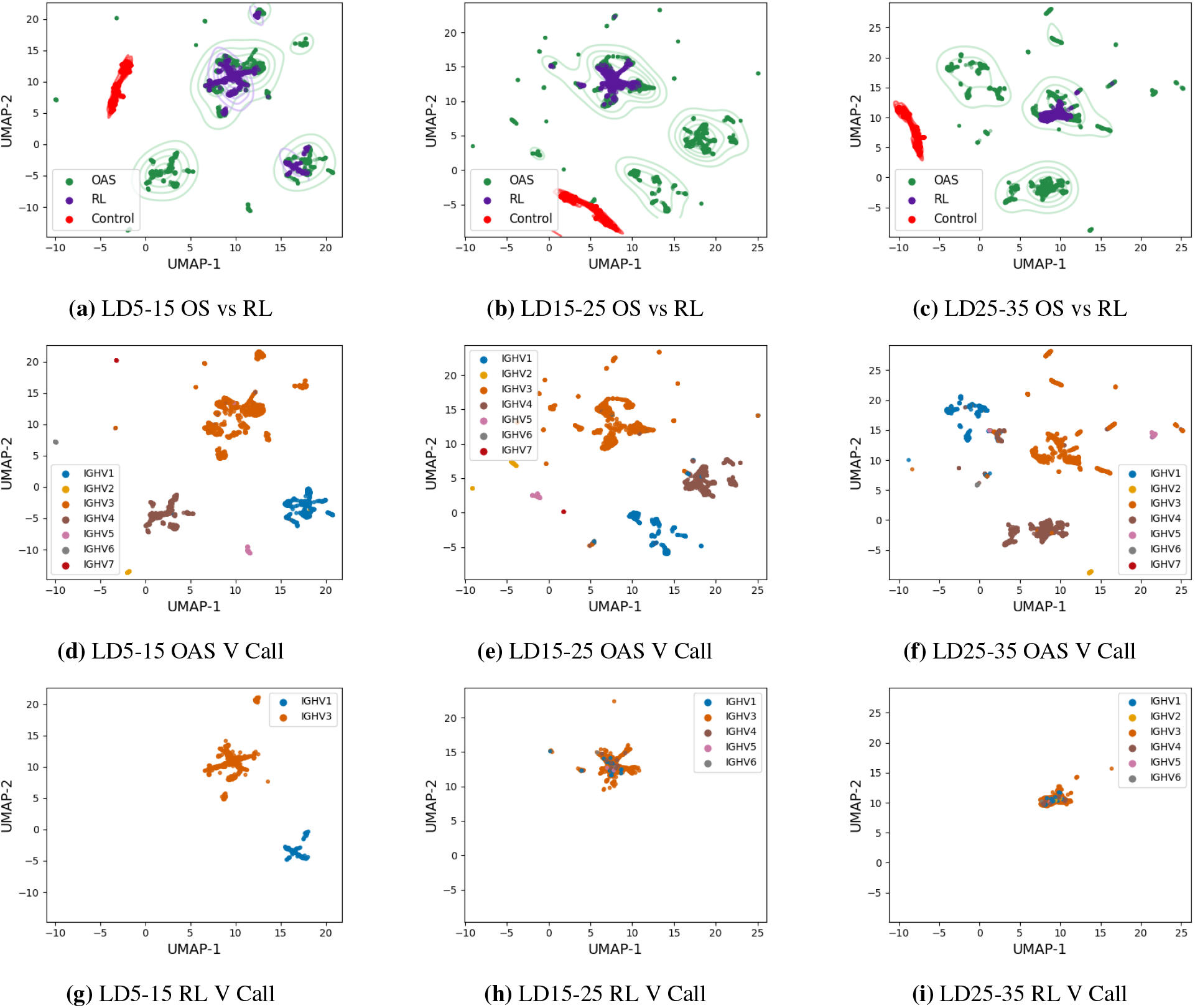
UMAP of AntiBERTy sequence embeddings for natural OAS and RL generated sequences from (**a**) LD5-15, LD15-25, (**c**) LD25-35. Contour lines represent the density of plotted embeddings, while the control comprises ribosome sequences. (**d**-**f**) UMAP OAS embeddings color-coded by V gene family. (**g**-**i**) UMAP RL embeddings color-coded by V gene family.

Across all LD bins, RL-generated antibodies occupy OAS sequence embedding clusters, indicating semantic consistency with naturally occurring antibodies. We also observe isolated clusters composed exclusively of OAS sequences, not populated by RL-generated antibodies.

We hypothesize that these isolated clusters correspond to germline families not explored by GermRL-trained policies. To test this, we color-coded UMAP projections by V (**Fig. 8**) and J (**Appx. G.1**) gene family for both RL and OAS antibodies. As previously observed [48], AntiBERTy embeddings primarily cluster by V gene identity, so RL and OAS sequences group together through shared V gene families. ProGen2-RL’s preference for IGHV3-derived sequences does not substantially populate clusters associated with other V gene families. Consequently, the isolated OAS clusters reflect the limited germline exploration induced by the bounded RL objective. This limitation is further shown in **Appx. F** through top-*k* cumulative density plots and presents an opportunity for future work to better capture the diversity of naturally occurring antibodies at elevated LD under the bound threshold objective.

### 6.5 RL-generated antibodies preserve CDR3 length, liability motif burden, stability and hydrophobicity profiles relative to natural antibodies

The heavy CDR3 region is highly variable and central to antigen specificity [49]. Both base ProGen2-OAS and ProGen2-RL preferentially generate heavy-chain antibodies (**Appx. E**). Although CDR3 residues are excluded from the LD reward, we examine whether RL training affects CDR3 length by comparing its distribution between RL-generated and OAS antibodies (**Fig. 9a**).

**Figure 9:**
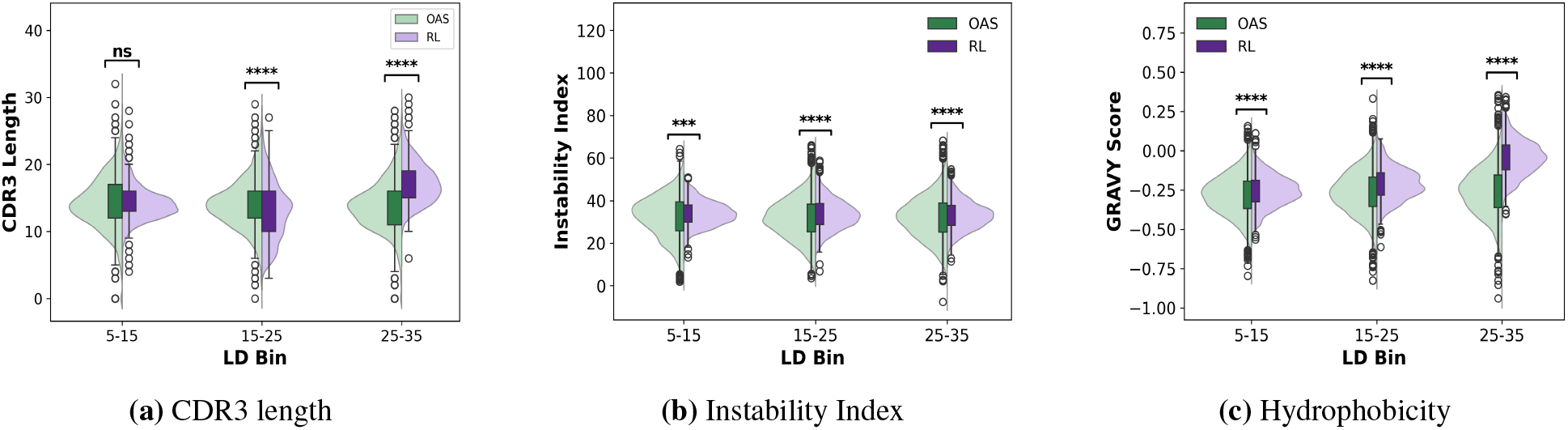
**(a)** CDR3 length, **(b)** instability index and **(c)** hydrophobicity GRAVY score between RL-generated and OAS sequences.

At LD 5-15, ProGen2-RL maintains CDR3 length distributions similar to natural antibodies, which is also consistent with base ProGen2-OAS sequences (**Appx. H.4a**). However, at LD 15-25, RL-generated antibodies exhibit slightly shorter CDR3 lengths, while at LD 25-35 the lengths are on average 3 to 4 residues longer. These results suggest that, at low LD, RL training does not substantially distort CDR3 length relative to natural antibodies and the pre-trained model. The modest shift at intermediate and high LD is consistent with our findings of different generative strategies explored by the policy.

Developability is a critical consideration in therapeutic antibody design. To assess whether RL training alters developability-associated properties, we compute instability indices [50] and GRAVY scores [51], the average hydropathy values of residues in a protein, for each sequence using the ProteinAnalysis module from Biopython [52] (**Fig. 9b, Fig. 9c**).

RL-generated antibodies show instability indices comparable to OAS antibodies (**Appx. 9b**) and consistent with base ProGen2-OAS generated sequences (**Appx. H.4b**), remaining below 40 and consistent with stable proteins. At LD 5-15 and LD 15-25, RL-generated antibodies exhibit similarly negative GRAVY scores to OAS antibodies and that of base ProGen2-OAS antibodies, indicative of maintained solubility of RL-generated antibodies in these LD bins (**Fig. 9c**), (**Fig. H.4c**). In contrast, at LD 25-35, RL-generated antibodies display higher GRAVY scores, shifting closer to zero relative to OAS antibodies and indicating lower solubility. Overall, RL training largely preserves key biophysical properties, while the increased hydrophobicity at higher LD reflects a trade-off associated with generating RL antibodies under higher LD objectives.

Finally, we examine whether GermRL increases the liability motif burden, or presence of residue patterns that cause unwanted modifications to antibodies, of RL-generated sequences. We compute the fraction of sequences containing liability-associated motifs across six liability categories (asp isomerization, asp-pro, missing cys, deamidation, extra cys, n-linked glycosylation), for both RL-generated and OAS sequences in each LD bin (**Fig. 10**).

**Figure 10:**
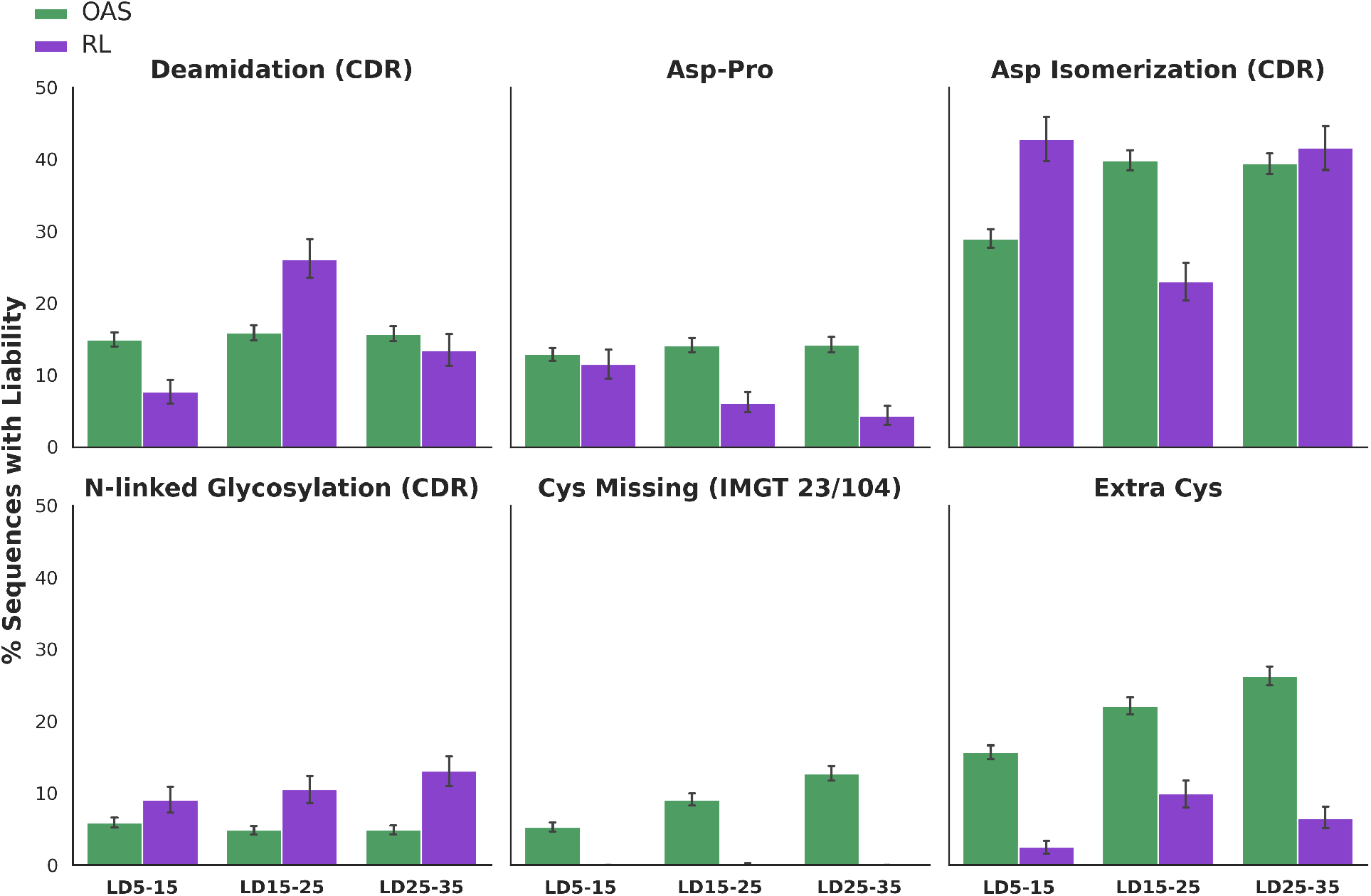
Percent of sequences containing liability motifs between RL generated and natural OAS sequences. Asp isomerization causes spontaneous change in protein backbone, Asp-Pro can result in peptide bond fragmentation, deamidation refers to the chemical degradation process that modifies residue side chains and N-linked glycosylation causes high heterogeneity. Asp isomerization, deamidation and N-linked glycosylation liabilities are searched for in each of the three CDR regions.

For both RL and OAS sequences, asp isomerization is the most frequent liability as it represents a common spontaneous change to the residue structural arrangement that affects protein stability and function. In vivo, this liability may be inconsequential given short protein lifetimes and quality control from the body through enzymes like PIMT [53]. However, for a therapeutic in storage, this liability can change the potency of the drug.

ProGen2-RL generates fewer sequences with extra cysteines. Similar to base ProGen2-OAS (**Appx. H.5**), RL-generated sequences do not exhibit missing cysteines at IMGT positions 23 and 104 compared to OAS. Liability burden is otherwise comparable between RL-generated and OAS sequences. These results indicate that RL training does not promote sequence liabilities, suggesting that the generated antibodies retain favorable developability properties relative to natural antibodies.

## 7 Discussion

Antibody therapeutic design requires sampling rare mutations to explore novel binding sites. However, antibody language models favor germline residues, limiting the diversity of generated sequences and their utility for antibody engineering. Our results demonstrate that RL can effectively steer autoregressive antibody language models away from the germline bias inherited during pre-training on large antibody repertoire datasets.

We began with default GRPO and introduced two modifications to stabilize RL optimization under challenging LD objectives, where policies would otherwise exploit reward shortcuts. Using modified GRPO within GermRL, we trained a series of ProGen2-RL policies with consistent one-shot generation success to produce antibodies far from germline while maintaining predicted foldability. GermRL achieves pass@1 scores of 0.992 and 0.950 for antibodies containing at least 5 and 35 mutations from germline (lowest and highest mutation threshold tested), respectively, compared to 0.398 and 0.034 for the base language model. We also introduced a bounded LD objective that produces policies capable of generating within specified LD ranges.

To assess knowledge gained during RL, we compared key biophysical properties between RL-generated and natural OAS antibodies. RL training induces systematic shifts in antibody composition relative to OAS that exceed the deviations introduced during pre-training. These shifts are most pronounced in CDR regions, which show enhanced mutational enrichment and altered amino acid preferences, while framework regions remain relatively conserved at lower LD. At the same time, RL-generated antibodies largely preserve developability-associated properties, including CDR length distributions, stability, hydrophobicity, and overall liability burden. Together, these findings suggest that RL-based generative models can expand exploration of antibody sequence space while retaining many characteristics of biologically plausible antibodies.

GermRL successfully generates antibodies further from germline, yet RL training at the bounded threshold objective significantly reduces the diversity of germline V gene calls represented in generated sequences, indicating that optimization toward LD-based objectives can unintentionally collapse exploration onto a narrow subset of germline families. Future work may address this through improved reward design. In addition, while ESMFold enables efficient pLDDT evaluation during training, more accurate folding oracles such as AlphaFold 3 [54] or Protenix [55] could provide stronger structural feedback.

More broadly, these findings support RL as a mechanism for controlled navigation of antibody sequence space. GermRL provides a practical approach for reducing the germline bias and diversifying mutations sampled from AbLMs in antibody engineering workflows such as binder design, infilling or fitness prediction. The framework may also serve as a foundation for designing RL objectives tailored to specific therapeutic challenges, while offering insight into how autoregressive antibody language models reorganize learned biological representations during RL optimization.

## Code Availability

GermRL training and inference scripts as well as trained ProGen2-RL policy weights are available at https://github.com/Graylab/GermRL.

## Appendix

### A Methods

#### A.1 Antibody levenshtein distance (LD) from germline

Levenshtein distance (LD) quantifies the divergence of an antibody sequence from its germline [56]. LD measures the minimum number of single-character edits needed to transform one sequence into another. In the antibody context, it can represent the number of mutations required to revert a sequence to its germline. To compute LD from germline, we identify V and J calls using ANARCI [57]. We then align the sequence separately to each call, truncate unaligned portions, and apply the Levenshtein algorithm to each region [56]. The final LD is the sum of the V and J region distances. Higher LD values indicate greater divergence from germline; LD = 0 indicates identity with germline. Heavy chain germline structure formally consists of V, D, and J regions. However, we excluded the D region from LD computation due to its high intrinsic variability.

#### A.2 Reward function

The core reward function for the single LD threshold objective has three components: an LD reward based on germline distance, a foldability reward based on ESMFold pLDDT, and penalties for edge cases. The bounded reward function extends the LD reward with an upper LD threshold (**Appx. F**).

##### A.2.1 Levenshtein distance (LD) reward

For each generated antibody, we first compute its LD from germline. If the policy generates a sequence with no identifiable germline, we assign an LD of −10. Given the computed LD, *S*_LD_, we apply the LD reward function defined in **Eq. 1** and shown in **Fig. A.1**.

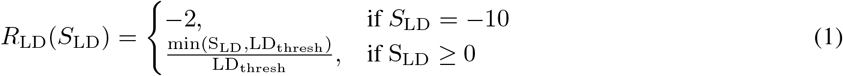

**Figure A.1:**
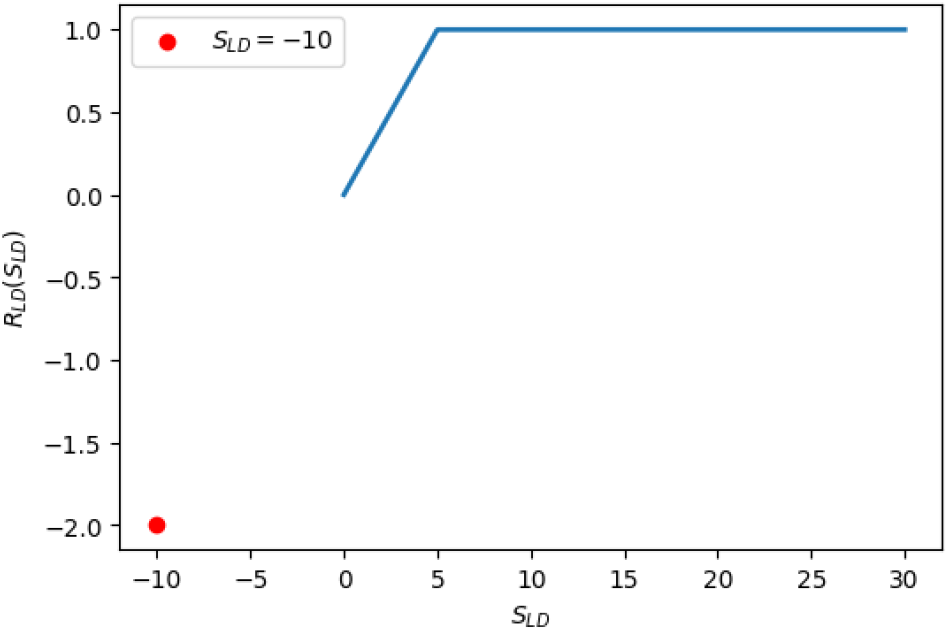
LD Reward where LD_thresh_ = 5. *S*_LD_ is the computed LD of a generated antibody sequence.

The LD reward assigns a maximum of 1 to sequences with LD ≥LD_thresh_, where LD_thresh_ is the minimum LD the policy should generate. The linear region before LD_thresh_ provides a steady reward signal guiding the policy toward the threshold. Once the threshold is reached, the reward is flat, where any sequence with LD≥ LD_thresh_ receives the same maximum reward. The penalty for sequences with no identifiable germline (LD = −10) is −2, twice the maximum reward, to strongly discourage actions that abandon the model’s pre-training entirely.

##### A.2.2 Levenshtein distance (LD) reward bounded

For the bounded LD threshold objective, we add both an upper threshold, 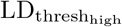, and a lower threshold, 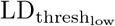 , to the reward function, shown in **Eq. 2**. Sequences within the desired LD range receive the maximum reward of 3, scaled higher than in the single-threshold objective. A buffer zone of 10 LD above 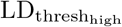 applies a linearly decreasing reward as *S*_LD_ exceeds the upper bound, discouraging the policy from exploring beyond the target region. Once 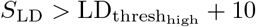, a flat penalty is applied.

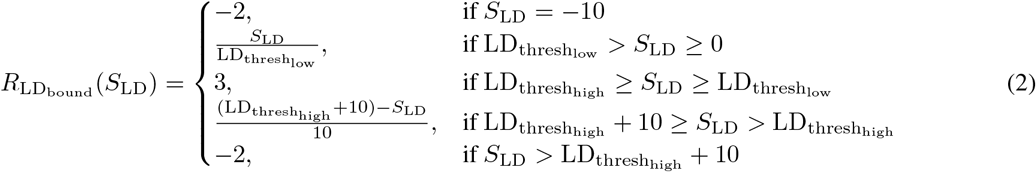

**Figure A.2:**
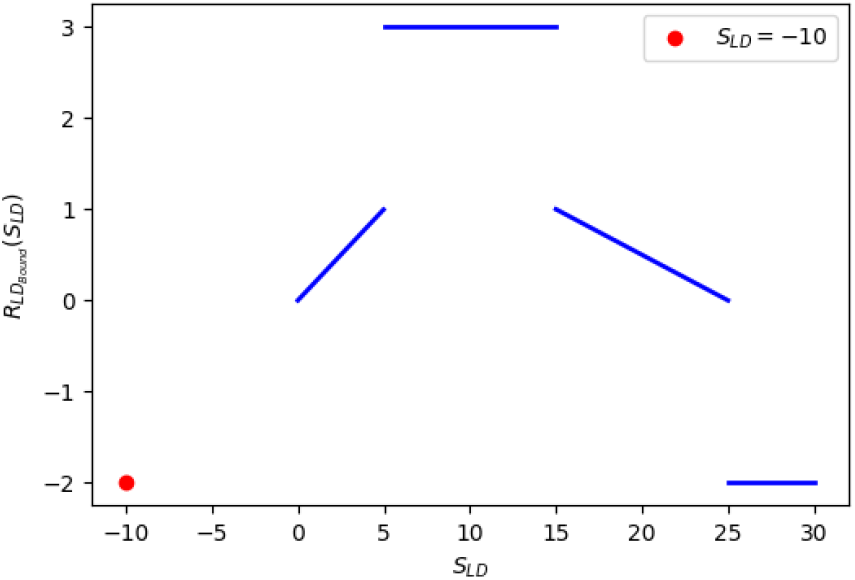
LD bounded reward where 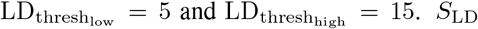 is the computed LD of a generated antibody sequence.

##### A.2.3 Foldability reward

While the previous reward function pushes the policy towards higher LD generations, the foldability reward maintains the structural integrity of the sequence. Each time a sequence is generated by the policy during training, we use ESMFold to compute the mean pLDDT, where pLDDT is a per-residue measure of confidence by structural protein AI models assigned to each amino acid position after folding a sequence. A higher ESMFold pLDDT value (maximum of 1) is a better indicator of the generated sequence and accompanying predicted structure looking similar to the natural distribution of proteins ESMFold was trained on, providing plausibility to the sequence.

Compared with other protein sequence to structure oracles, we chose ESMfold for its speed. This is a design choice for this framework to accelerate the training process. The ESMFold pLDDT of a generated antibody is defined as *S*_P_ and is used in the following reward in **Eq. 3**, graphed in **Fig. A.3**.

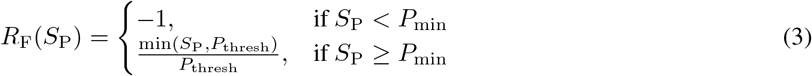

*P*_min_ and *P*_thresh_ represent the minimum pLDDT before a penalty is introduced and the threshold pLDDT where the policy should be generating sequences above to receive the maximum reward, respectively. Above *P*_min_, the reward function is linear until *P*_thresh_ is reached. However, unlike LD, base ProGen2-OAS can consistently generate sequences with good pLDDT at the start of training, as it was pre-trained on foldable antibody sequences. Therefore, the difference between *P*_thresh_ and *P*_min_ is small. Mainly, this reward function discourages the policy from exploring actions with low foldability.

**Figure A.3:**
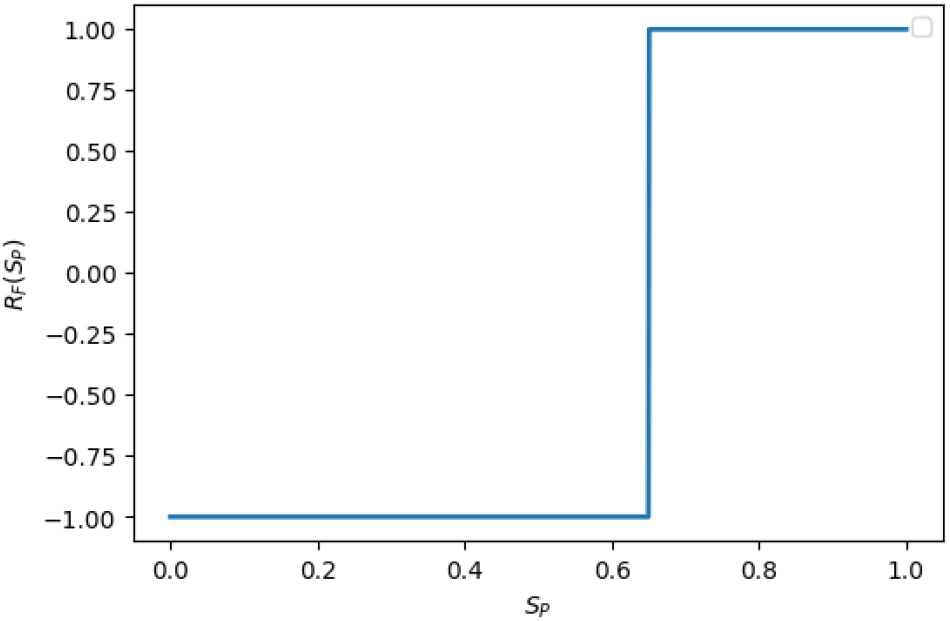
Foldability reward where *P*_thresh_ = 0.7 and *P*_min_ = 0.65. *S*_P_ is the computed ESM pLDDT of a generated antibody sequence.

##### A.2.4 Edge case penalties

During training, the policy can explore unfavorable action spaces resulting in nonsensical generations. For these cases, a flat penalty of -2 is applied. There are three edge cases taken into account. If the antibody sequence:

- Is a very short sequence, specifically if it has < 100 residues.
- Has non amino acid tokens between the start and end token.
- Has a non human closest germline, determined using ANARCI.

##### A.2.5 Reward germline trajectory (RGT) reward in bounded LD objective

To maintain base ProGen2-OAS germline call diversity throughout RL training, we introduce another reward component, named Reward Germline Trajectory (RGT), that adds an incentive for V call diversity. The RGT reward can be added or removed during RL training. While training with the bounded LD objective, we notice that some policies favor certain V calls, while J calls maintain a similar level of diversity observed in base ProGen2-OAS. To discourage this behavior, RGT diminishes the total reward (the sum of the LD and fold reward components) of generated sequences with a germline V call seen multiple times during the training epoch. Each time a trajectory is generated during training, we use ANARCI [57] to find the germline V call and keep track of the fraction of total antibodies generated in the epoch that belong to the call. Using this value, scaled by weight factor *κ*, we compute a reduce factor, 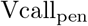, shown in **Eq. 4** where pop_V_ is represents the number of times the generated antibody sequence’s V call was previously generated and 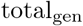 is the total number of generations in a training epoch.

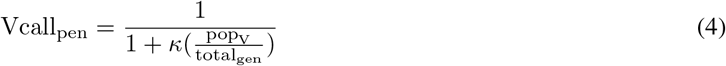

When RGT is applied, the sum of the reward components (LD and Fold), is scaled by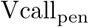. Therefore, a generated sequence with good LD and pLDDT can receive a poor reward if the antibody belongs to a germline call that makes up the majority of generations in the epoch.

#### A.3 GRPO introduction

RL training is commonly formulated as a Markov decision process (MDP), in which a policy begins from an initial state and sequentially selects actions. After each action, the environment returns a reward and transitions to a new state [58]. RL has become an important component of large language model (LLM) training for natural language processing tasks [32]. Unlike pre-training, which optimizes models to reproduce patterns from large text corpora, RL enables models to learn through iterative exploration and feedback. This paradigm has improved performance on complex reasoning tasks, including coding and mathematics, and supports safety alignment and tool use in agentic workflows [33], [34].

Autoregressive (AR) language models generate tokens sequentially from left to right, using existing context to predict the next token. The probability of generating a sequence is expressed as the joint probability 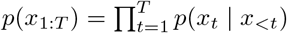, where *T* is the sequence length and each token *x*_*t*_ is conditioned on all prior tokens *x*_*<t*_. This can be framed as an RL episode where *x*_*t*_ is the action (chosen token) and *x*_*<t*_ is the state (prior context). Once a token is selected, the state updates to *x*_*<t*+1_, and the process continues until the full sequence is generated. This mirrors an MDP in which the AR language model serves as the policy, choosing tokens (actions) given the existing context (state).

**Figure A.4:**
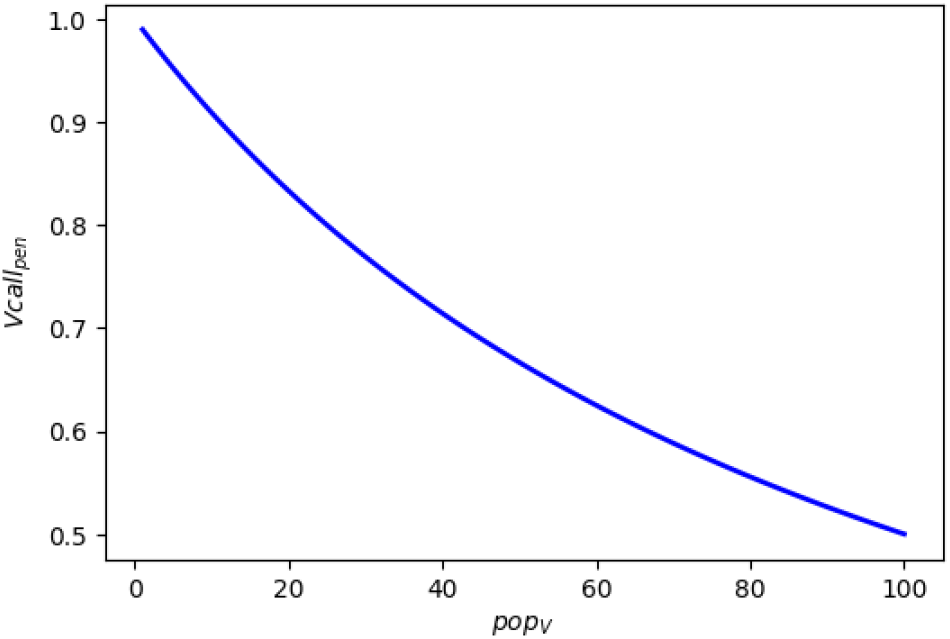
Reduce factor Vcall_pen_ that is applied to the sum of LD and pLDDT reward during application of RGT for increasing popularity of the generated antibodies’ V call, pop_V_, when *κ* = 1 and total_gen_ = 100.

To alleviate the germline bias, we use Group Relative Policy Optimization (GRPO), a policy gradient algorithm that directly optimizes policy parameters through gradient ascent on rewards [37]. GRPO samples multiple trajectories from the policy *π*_*θ*_ for the same input prompt, forming a batch. Each output is evaluated using a reward function, and the policy is updated toward higher-reward trajectories. The GRPO objective is 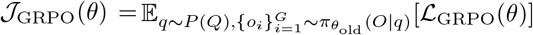, where *q* ∼ *P* (*Q*) denotes sampling a prompt *q* from the dataset *Q* and 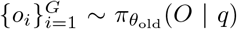 (*O* | *q*) represents the generation of *G* trajectories using the old policy 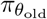 . During training, two versions of the policy are maintained: *π*_*θ*_, the currently updating policy, and 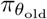, an older version that serves as the trust-region baseline. In default GRPO, after each generation step, *π*_*θ*_ updates and then synchronizes weights with 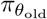, so that the old policy reflects recent updates at the start of the next step. ℒ_GRPO_(*θ*) is the GRPO loss defined in **Eq. 5**.

The probability ratio 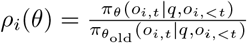 compares the current and old policy distributions over the chosen token *o*_*i,t*_ at position *t* in trajectory *i*, given prompt *q* and prior tokens *o*_*i,<t*_.

*ρ*_*i*_(*θ*) is scaled by the advantage, 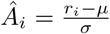 , a measure of how trajectory *i* performs relative to other trajectories in the generated batch and is defined by taking the z-score of all rewards in the batch, where *μ* is the mean and *σ* the standard deviation.

*ρ*_*i*_(*θ*) follows a clipped surrogate objective from PPO [36] to stabilize policy updates. To prevent large deviation from pre-training, the GRPO surrogate loss includes a KL penalty with weighting factor *β*, measuring KL divergence between *π*_*θ*_ and *π*_ref_ , a frozen copy of the pre-trained policy. The complete surrogate loss ℒ _GRPO_(*θ*) is shown in **Eq. 5**, averaging the clipped and scaled *ρ*_*i*_(*θ*) across all tokens in each trajectory and over all *G* trajectories in the batch [37].

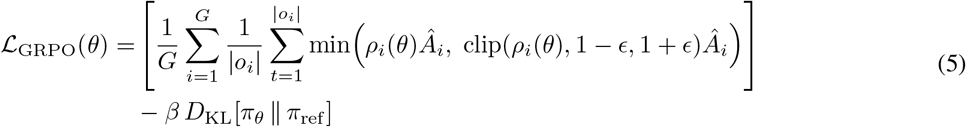

To maximize the GRPO objective function in optimizing the policy, we follow the outcome supervision RL training scheme described by Shao et al. where a single reward is assigned at the end of each output. The GRPO training algorithm is shown in **Alg. 1**.

##### Algorithm 1

GRPO algorithm adapted from Shao et al. [37]

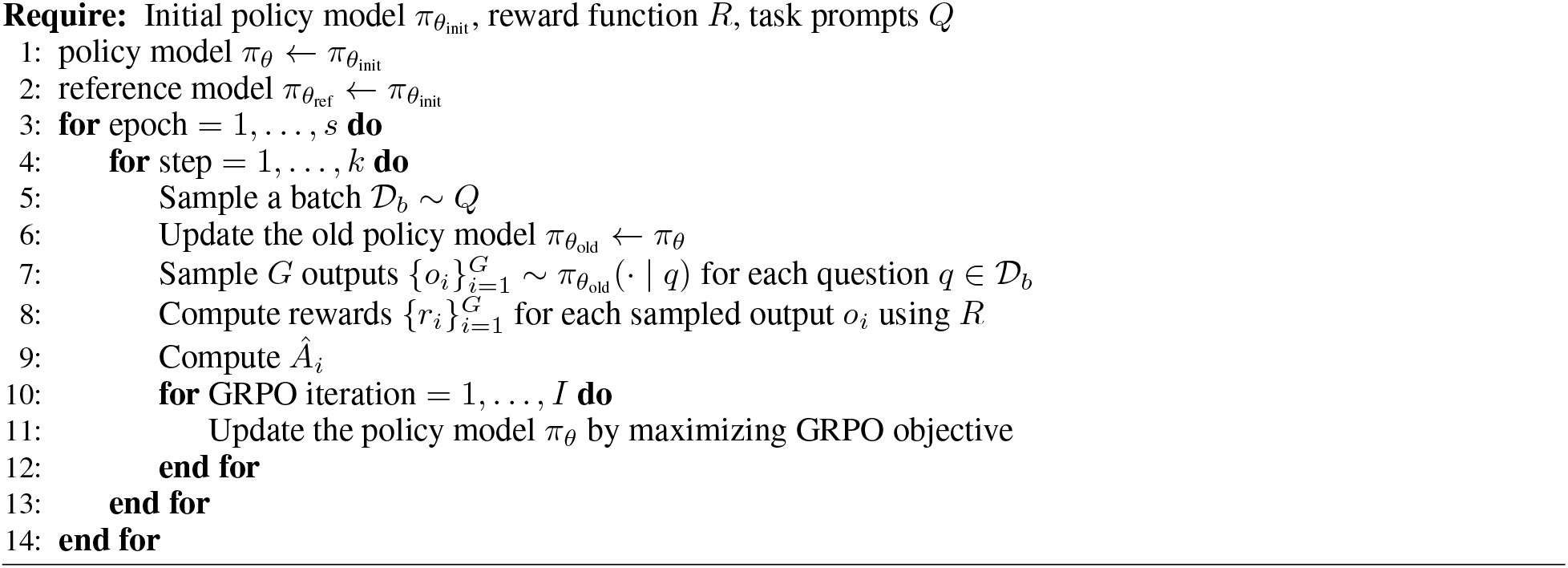

We note that in the original algorithm, the number of GRPO iterations *I* is referred to as *μ* [37]. When *I* = 1, GRPO is ran as an on-policy training regime, since the policy updating is the same as the policy sampling the trajectories. However, when *I >* 1, GRPO training follows an off-policy scheme as *π*_*θ*_ updates for multiple iterations while observing the same trajectories generated by 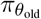 at the start of the training step [37],[59].

#### A.4 GRPO additions from previous work

We incorporate two additions to the base GRPO algorithm from prior work; we refer to the resulting configuration as “default GRPO.” First, we apply the clip higher strategy from dynamic sampling policy optimization (DAPO) [41]. This modification raises the upper clipping bound in ℒ_GRPO_(*θ*), increasing the probability that the updating policy can assign to beneficial but underexplored actions.

Second, following practical implementations of GRPO for LLM training [40], we add an entropy loss to encourage diverse generations. Our entropy loss, defined in **Eq. 6**, computes Shannon entropy over the distribution of all possible tokens *n* at each action step. This is first averaged across all actions within a trajectory, then averaged over the *G* trajectories in the batch.

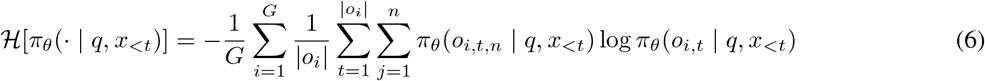

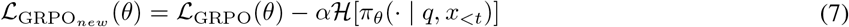

Here, *π*_*θ*_(*o*_*i,t,n*_| *q, x*_*<t*_) is the probability of each possible action taken by the policy, given the state, within the trajectory *i*. This probability is multiplied by the log probability, and this product is summed for all actions that make up the trajectory. We compute the final entropy by repeating this calculation across all trajectories in the batch *G* before taking the mean and negating the result. The entropy loss discourages overly deterministic action selection by maximizing the objective when the policy maintains high entropy, corresponding to a more distributed probability over possible actions at position *t* in a trajectory. Similarly to the KL penalty, we add a weighting term *α* to the entropy value before integrating it in the GRPO surrogate loss to obtain the updated GRPO loss shown in **Eq. 7**.

#### A.5 Custom GRPO modifications

Our GRPO modifications are written out in **Alg. 2** and detailed in this section.

##### Algorithm 2

Modified GRPO algorithm

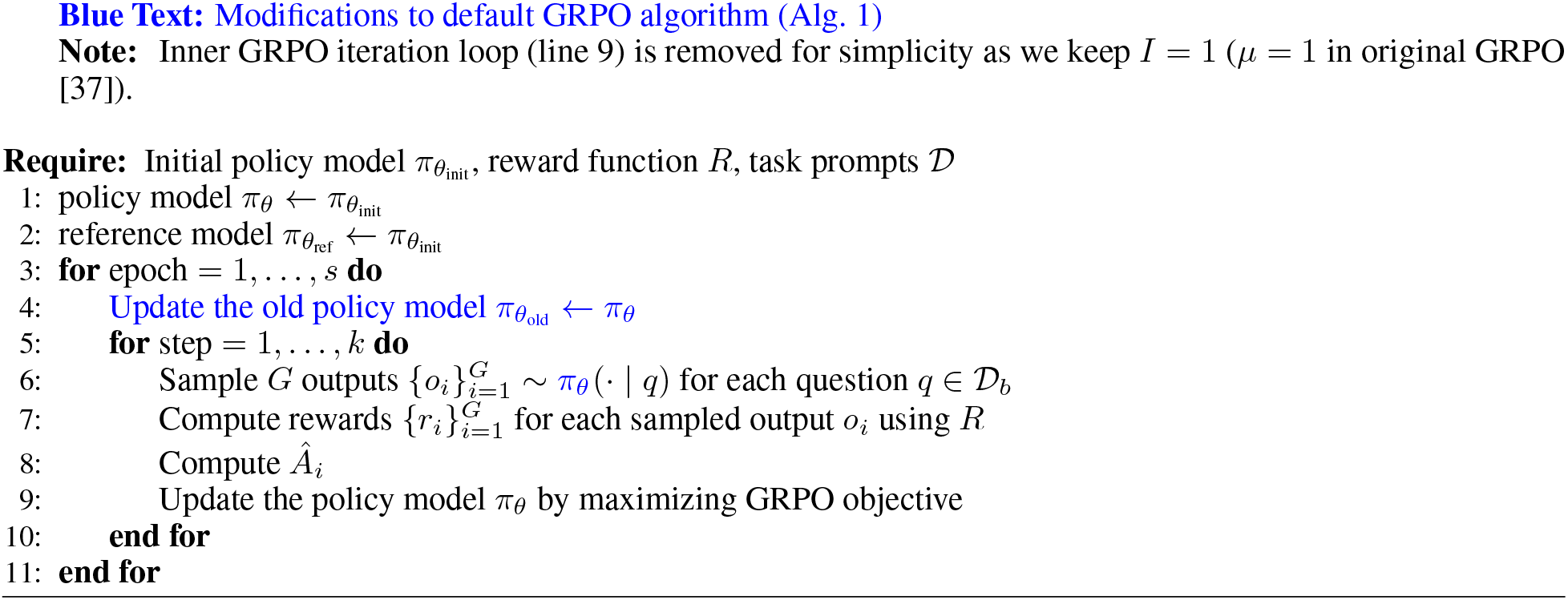

To improve policy training for our antibody application, we propose a pair of modifications to the default GRPO training algorithm. First, we set the old policy synchronization outside of the step loop, where trajectories are generated, and under the outer epoch loop. Now, multiple policy updates occur before the updating policy and old policy synchronize weights. We note that this modification alone was similarly described by Mroueh et al. [59] where they demonstrate an alternate off-policy training to that explored by Shao et al.

Mroueh et al. denote *v* as an update frequency parameter within *k* steps when 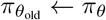 weight synchronization occurs. Using their notation in our algorithm, *v* = *k* such that updating the old policy weights occurs once per epoch when we iterate through all steps. Mroueh et al. define their off-policy GRPO scheme when *v >* 1 while *I* = 1. Therefore, synchronization of the old policy weights no longer occurs at every update step, while the current policy continues to be updated once per step. As a result, the policy used for sampling is no longer the same as the policy being updated.

Using the same notation, the off-policy regime described by Shao et al. would now be defined by *v* = 1 and *I >* 1. With their off-policy training scheme, Mroueh et al. demonstrate superior performance over the original from Shao et al. [59].

A primary motivation for Mroueh et al. is the reduced computational costs traditionally incurred through frequent weight synchronization of the old policy every step. Given the smaller scale of our antibody language model, we are not primarily concerned with expensive computation but rather introduce our set of modifications to ensure that the old policy provides a stricter baseline for future updates by limiting synchronization frequency with the newly updated policy.

Unlike Mroueh et al., we keep 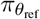 frozen during training, we do not use binary rewards (**Appx. A.2**) and, introduce a second key modification, where we generate trajectories from the current policy rather than the old policy. With this change, the updating policy and sampling policy remain the same, maintaining an on-policy RL scheme even as the old policy lags behind.

We maintain an on-policy scheme for two reasons. First, since the old policy remains close to the germline-biased initialization, using its generations in the probability ratio slows the opportunity to evaluate the higher-LD sequences discovered through recent policy updates. Second, the inclusion of this additional modification promotes stable training as, under LD35, RL training completely collapses while sequence identity remains elevated under the application of only the 1st modification (**Fig. A.5**).

Overall, we lag the old policy behind the updating policy and sample exclusively from the latter, allowing multiple rounds of trajectories from an already-updated policy to be observed before constructing a new trust region during synchronization.

Under this modified training scheme, high-reward trajectories identified early in training are less easily consolidated into the weights of the old policy, preventing them from prematurely defining the update baseline and driving further exploitation of these trajectories. We reason that these GRPO modifications better adapt the algorithm to our specific antibody optimization objective by keeping the policy robust to favoring successful actions discovered early in training under difficult objectives.

**Figure A.5:**
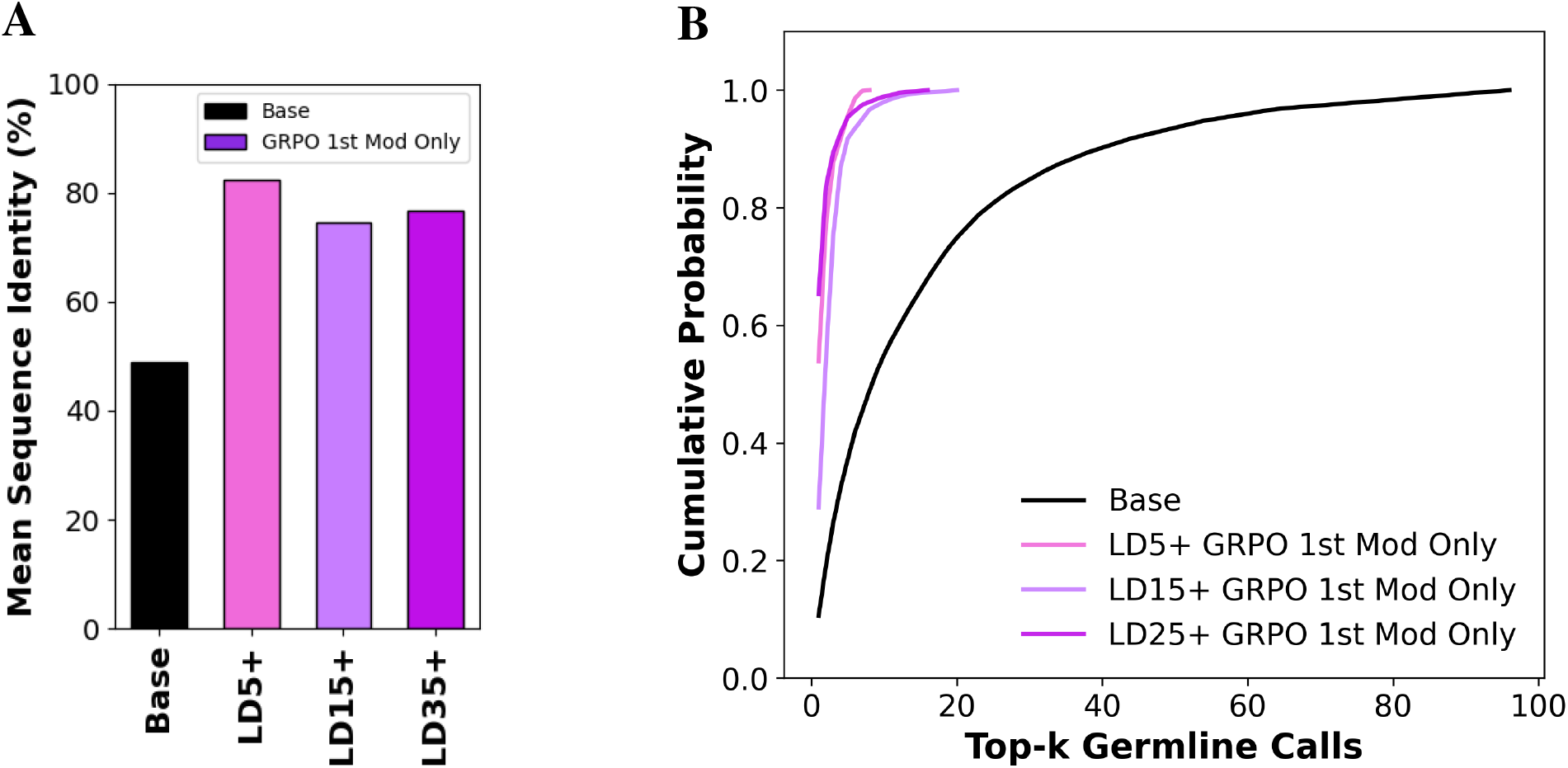
**(a)** Mean sequence similarity between each pair of antibodies from all base ProGen2-OAS generations and policy generations after RL training with modified GRPO using the 1st modification only. **(b)** Cumulative distribution of top-*k* germline V calls of generated sequences from base ProGen2-OAS and after RL training with modified GRPO using the 1st modification only.

#### A.6 RL training parameters

For the foldability reward we keep *P*_thresh_ to 0.7 and *P*_min_ at 0.65 (**Appx. A.2**). We run all single LD threshold training schemes for 80 epochs and bounded LD thresholds at 100 epochs. Each epoch contains 5 steps. For each step, we generate 12 trajectories through the policy. We set the number of update cycles, to *I* = 1 to follow the default on-policy GRPO methodology described by [37]. Therefore, for a batch of generated trajectories, there is only 1 policy update before old policy weight synchronization at the beginning of the next step. We use 0.1 as the clip higher parameter, as suggested in DAPO [41]. Furthermore, we use an entropy weight term of 0.03 while the KL divergence penalty weight term is 0.04. Finally, we use a learning rate of 0.00001 and ADAM optimizer for policy updates.

### B Generated antibody identifiable germline calls

**Figure B.1:**
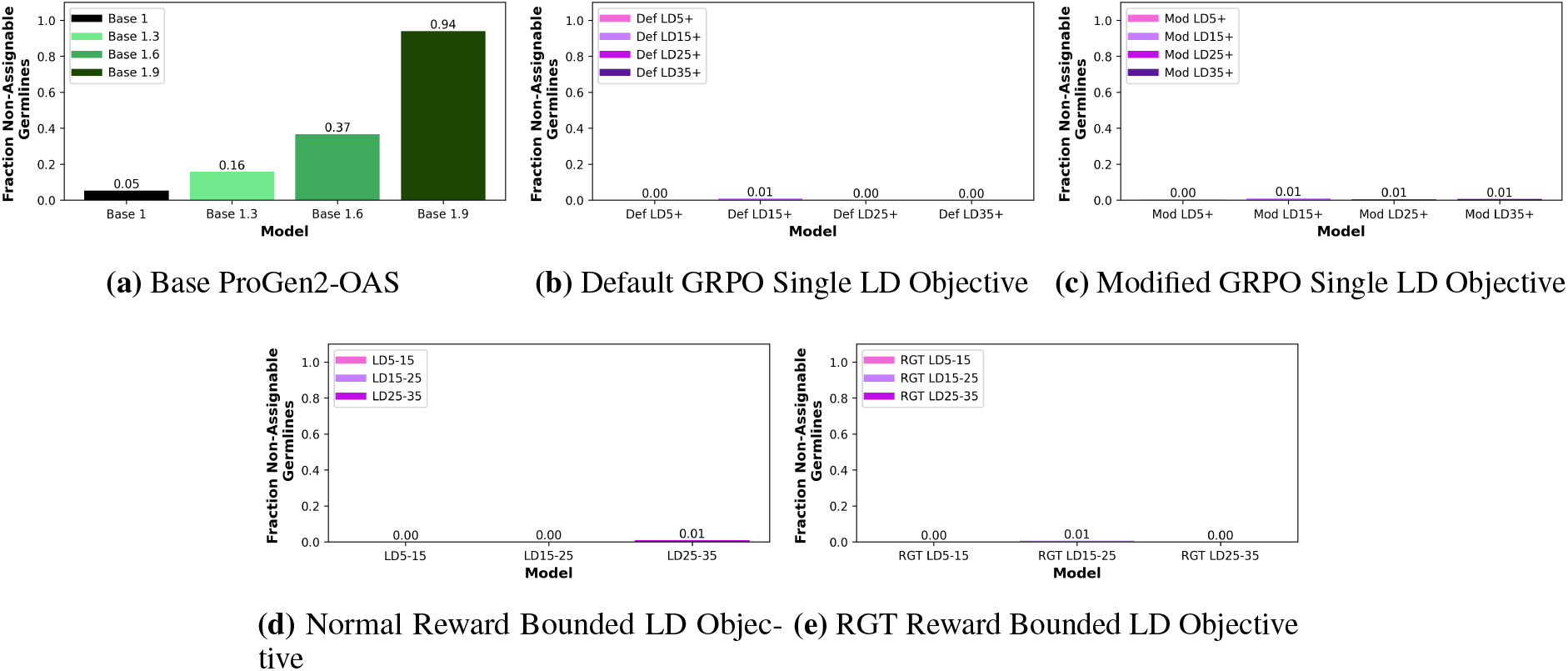
Fraction of generated antibodies from base ProGen2-OAS and ProGen2-RL policies that have a non-identifiable germline call.

### C RL training pass@k performance

To measure RL training performance, we calculate the mean pass@*k* across all steps for each epoch. Pass@*k* defines the probability where, given *k* generation attempts by the model, at least 1 is a success [60]. We use pass@1 which represents the probability that, given only one generation attempt by the model, it is a success. We define a successful antibody sequence generation as one that :

- Is > 100 residues
- Has only canonical amino acid tokens
- Has an identifiable human germline as the closest germline call
- Has LD *>*= LD_thresh_ (single LD objective) or *LD* within LD threshold bounds (bounded LD objective)
- Has *P >*= *P*_thresh_

### D Generated antibody sequence logo plots

We build logogplots of 1,000 antibody sequence generations for each RL trained policy with GermRL and from base ProGen2-OAS. We create the plots by first assigning IMGT numbering to each generated antibody sequence. Next, the sequence with the most numbered positions is set as the max length, and every other sequence is assigned filler positions to meet the same length. In the plot, a gray dash represents no residue at that numbered position. Through the logoplots, we demonstrate the truncation bias present in OAS that affects base ProGen2-OAS as well as the entropy collapse from training with default GRPO and alleviated with modified GRPO.

#### D.1 Base ProGen2-OAS

**Figure D.1:**
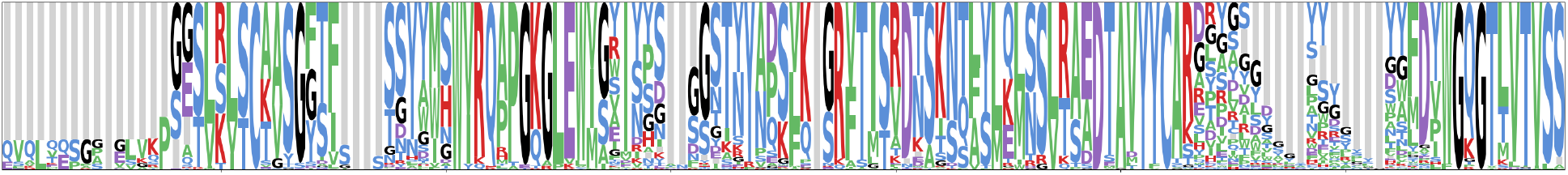
Logo plot from antibody sequence generations from base ProGen2-OAS

#### D.2 Default and modified GRPO single LD threshold

**Figure D.2:**
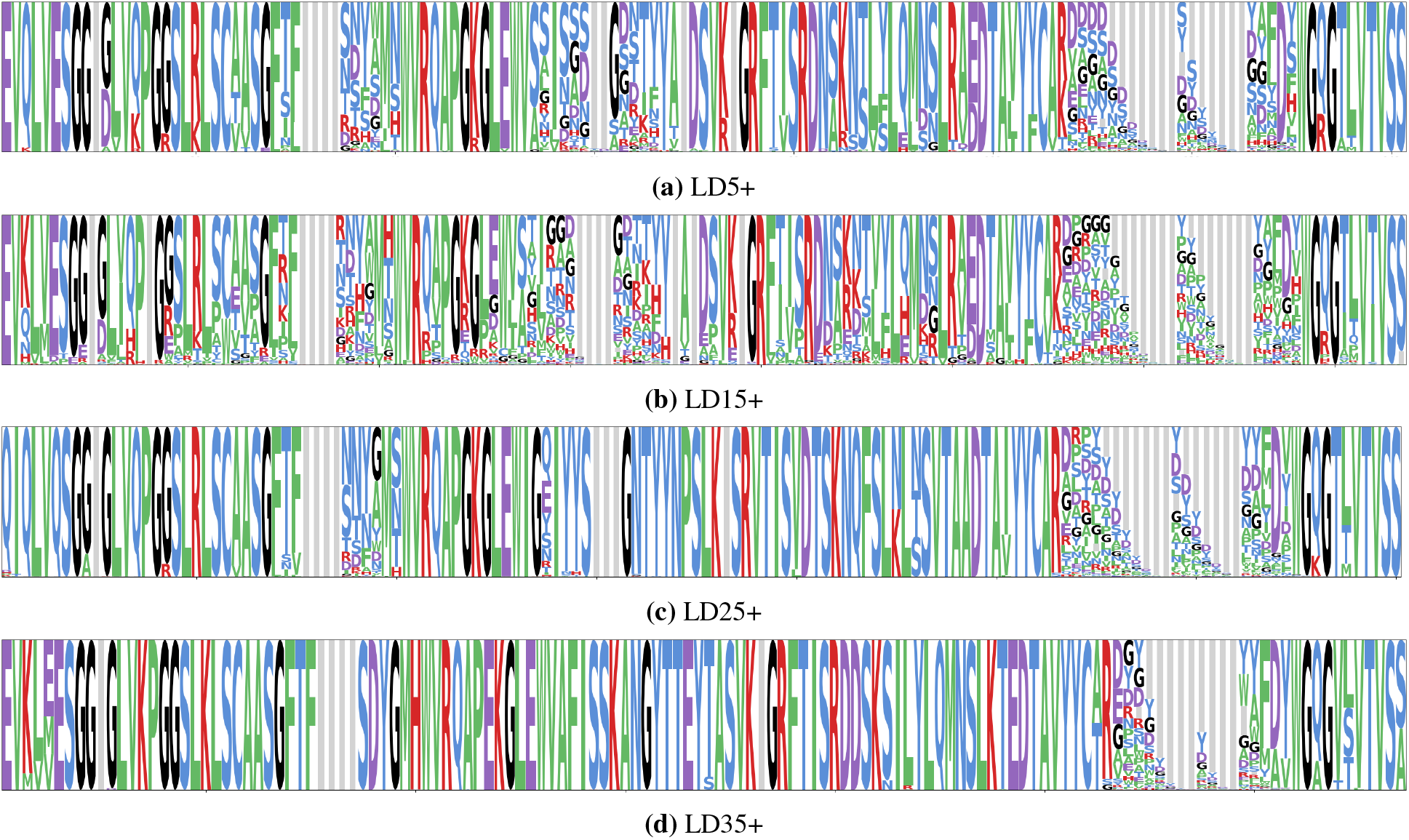
Single LD threshold RL trained with default GRPO policy logo plots.

**Figure D.3:**
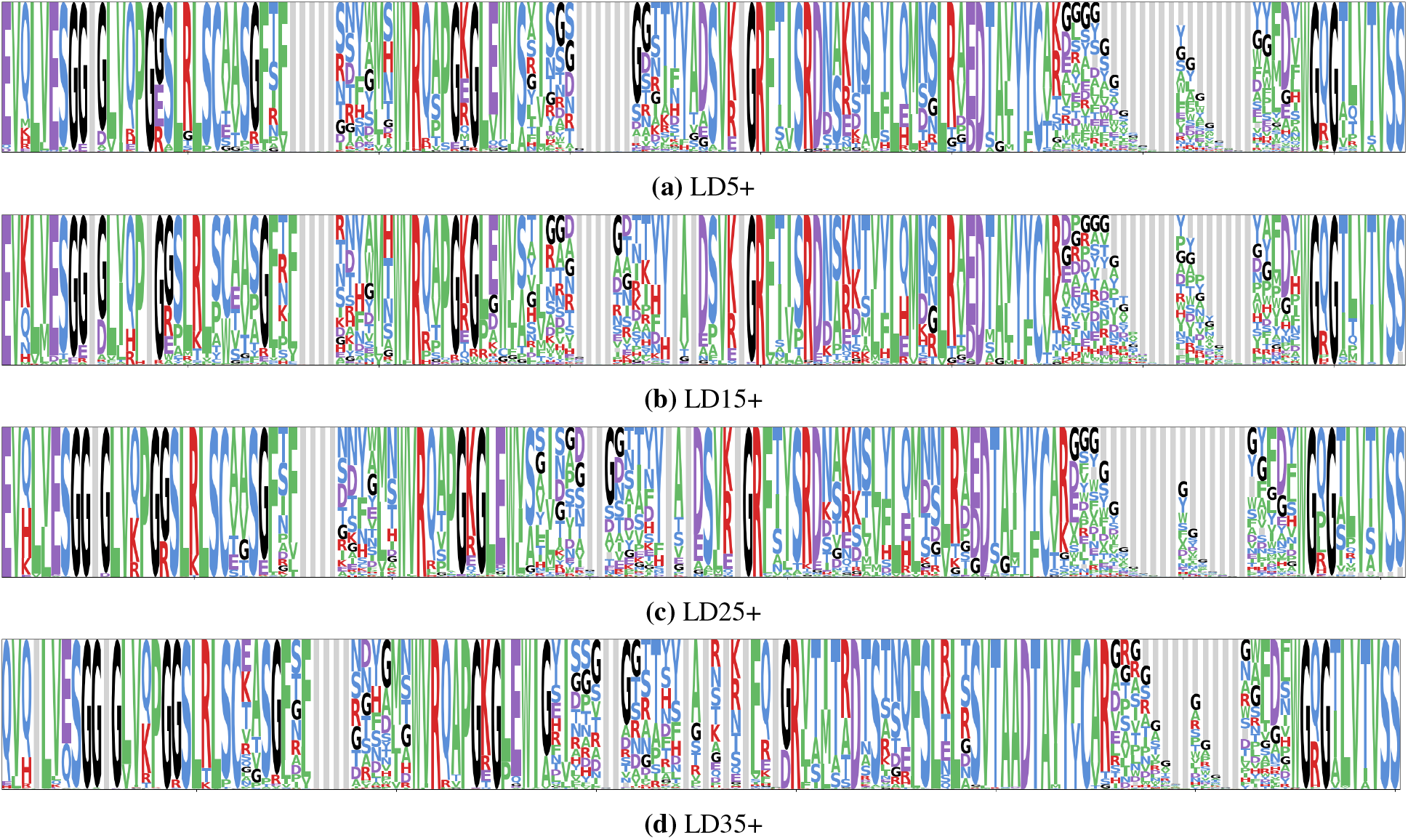
Single LD threshold RL trained with modified GRPO policy logo plots.

#### D.3 Normal and RGT reward bounded LD threshold

**Figure D.4:**
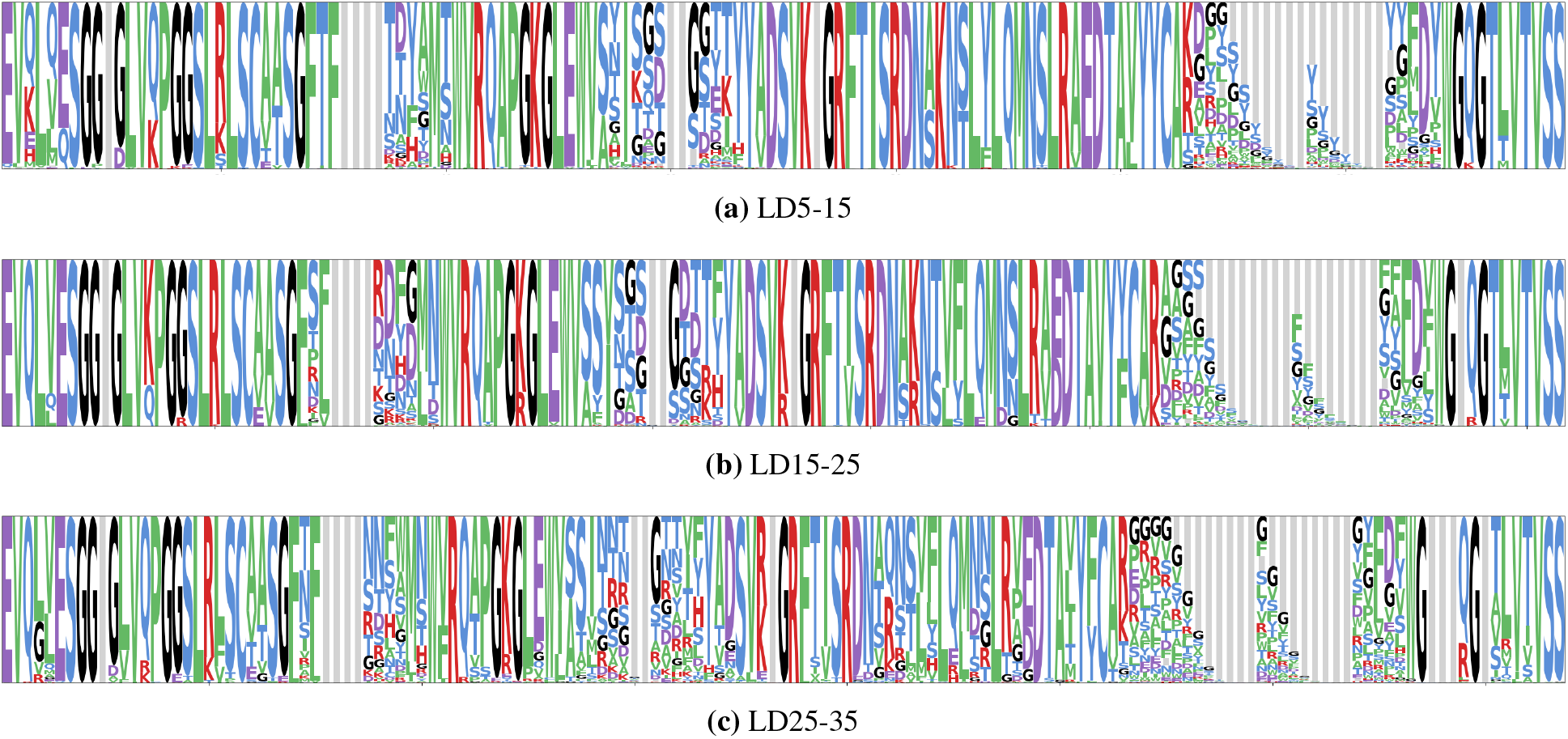
Bounded LD threshold RL trained with modified GRPO and normal reward policy logo plots.

**Figure D.5:**
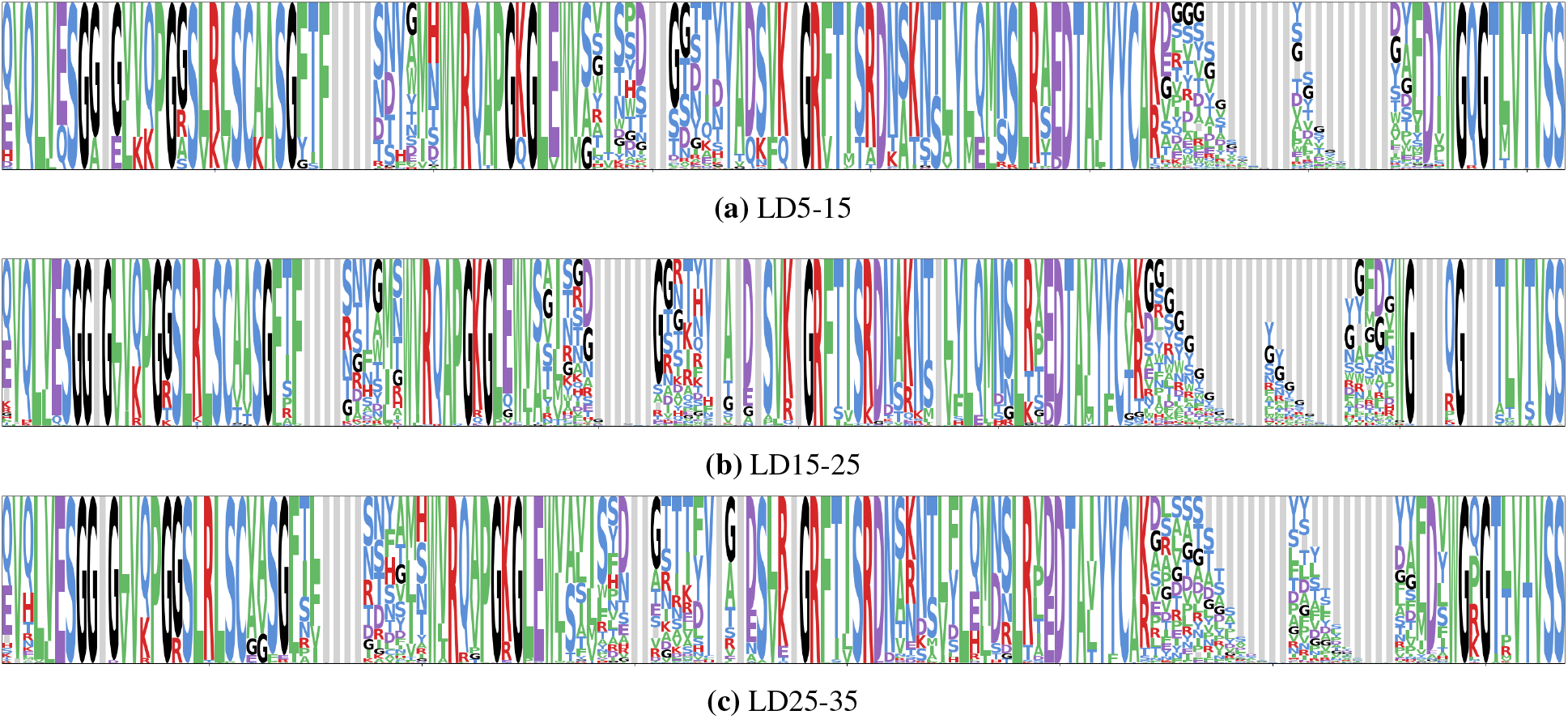
Bounded LD threshold RL trained policy with modified GRPO and RGT reward logo plots.

### E. Distribution of V and J germline calls across all RL trained policies and base Progen2-OAS

We build pieplots of germline V and J calls from the 1,000 antibody sequence generations for each RL trained policy and from base ProGen2-OAS. We assign calls through ANARCI [57]. Under single LD threshold RL training with default GRPO we demonstrate the policy entropy collapse, alleviated with modified GRPO. Although J call diversity remains similar, compared to the V call diversity generated with base ProGen2-OAS, we observe fewer unique V calls explored by all RL policies. Specifically, ProGen2-RL tends to favor IGHV3 family V calls, representing a limitation to address in future work.

#### E.1 Base ProGen2-OAS unique germline calls

**Figure E.1:**
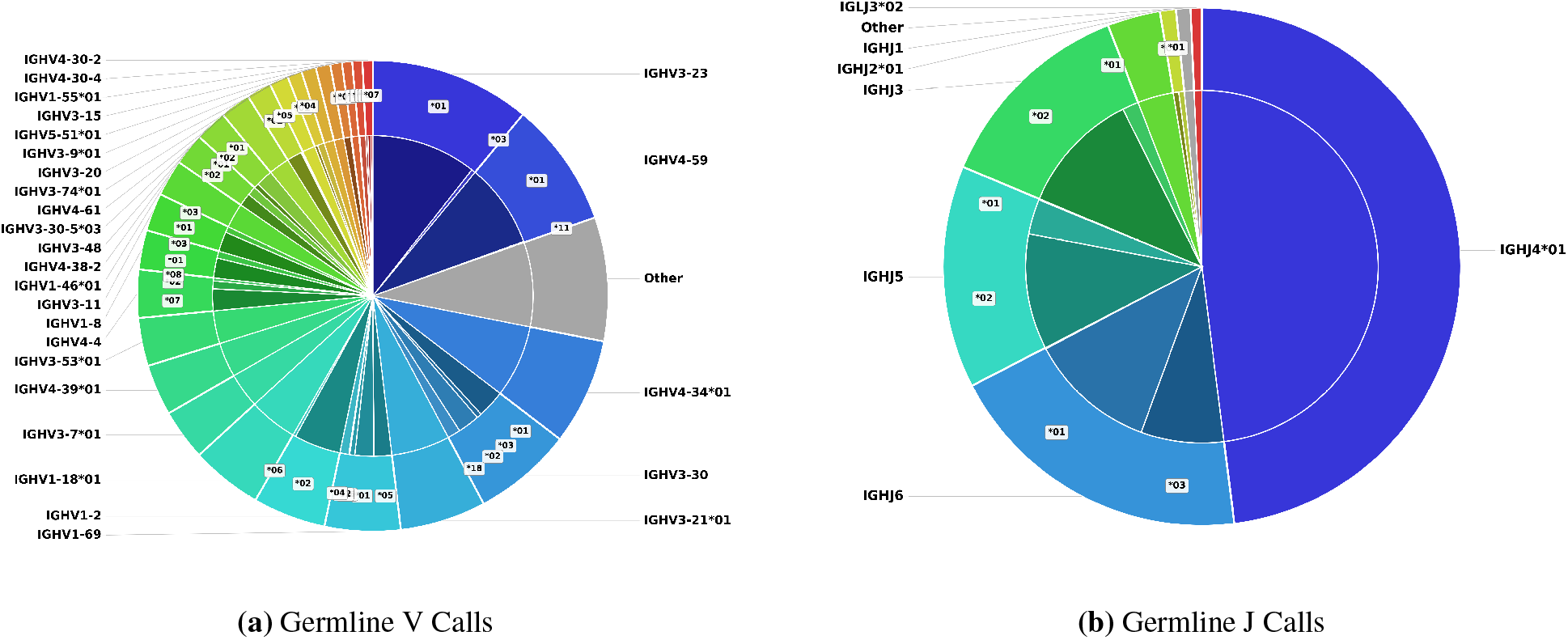
Distribution of unique germline V and J calls for Base Progen2-OAS at temperature 1.0

**Figure E.2:**
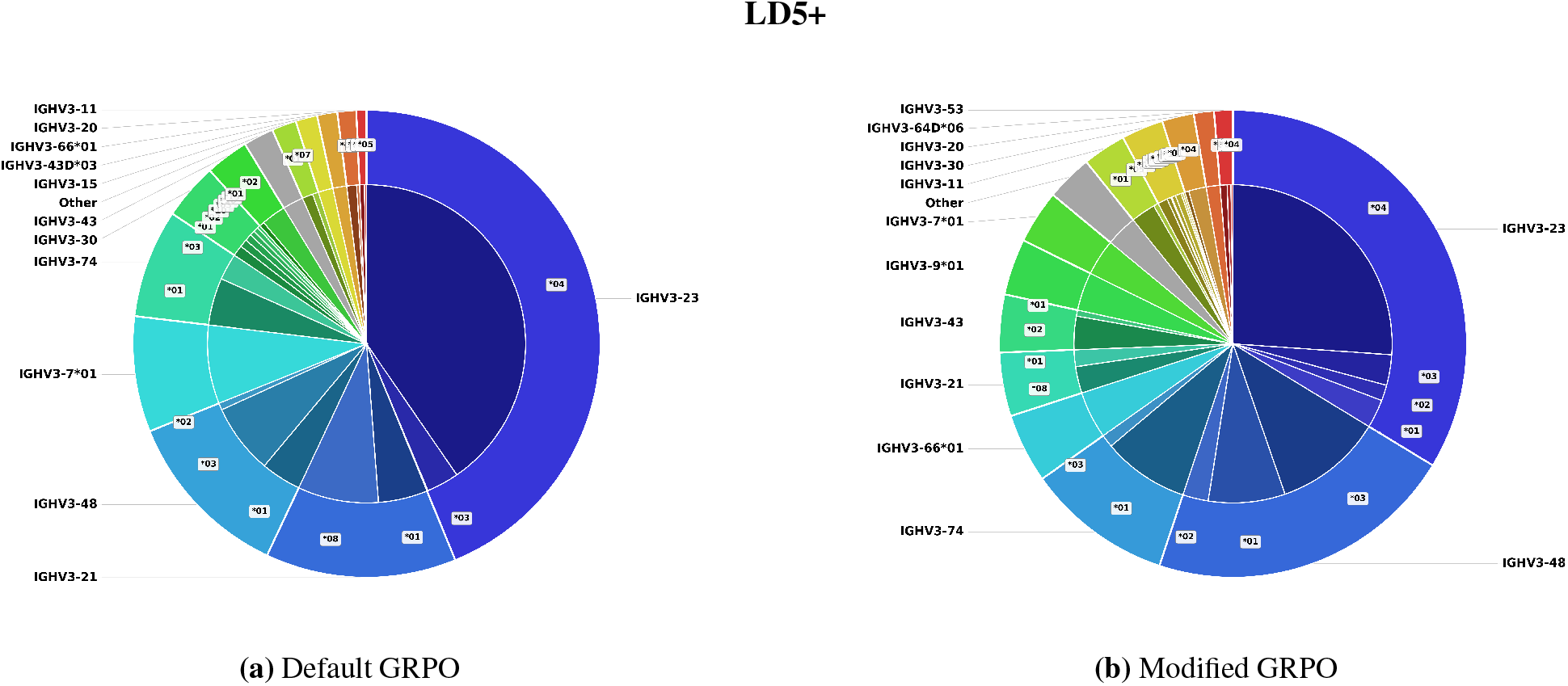
Distribution of unique germline V calls at LD5 threshold.

#### E.2 Single LD_Thresh_ modified and default GRPO training unique germline calls from sequence generations after RL training

##### E.2.1 Unique V calls

**Figure E.3:**
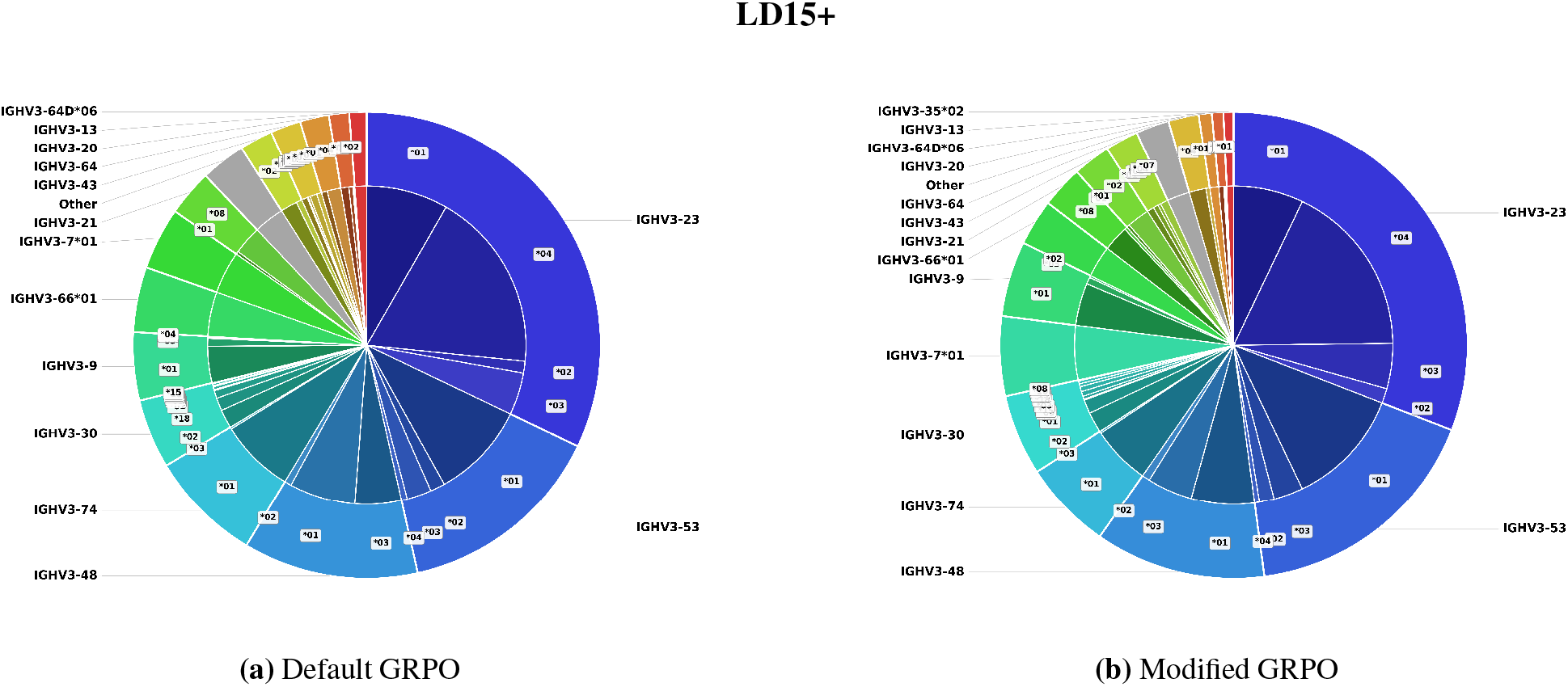
Distribution of unique germline V calls at LD15 threshold.

**Figure E.4:**
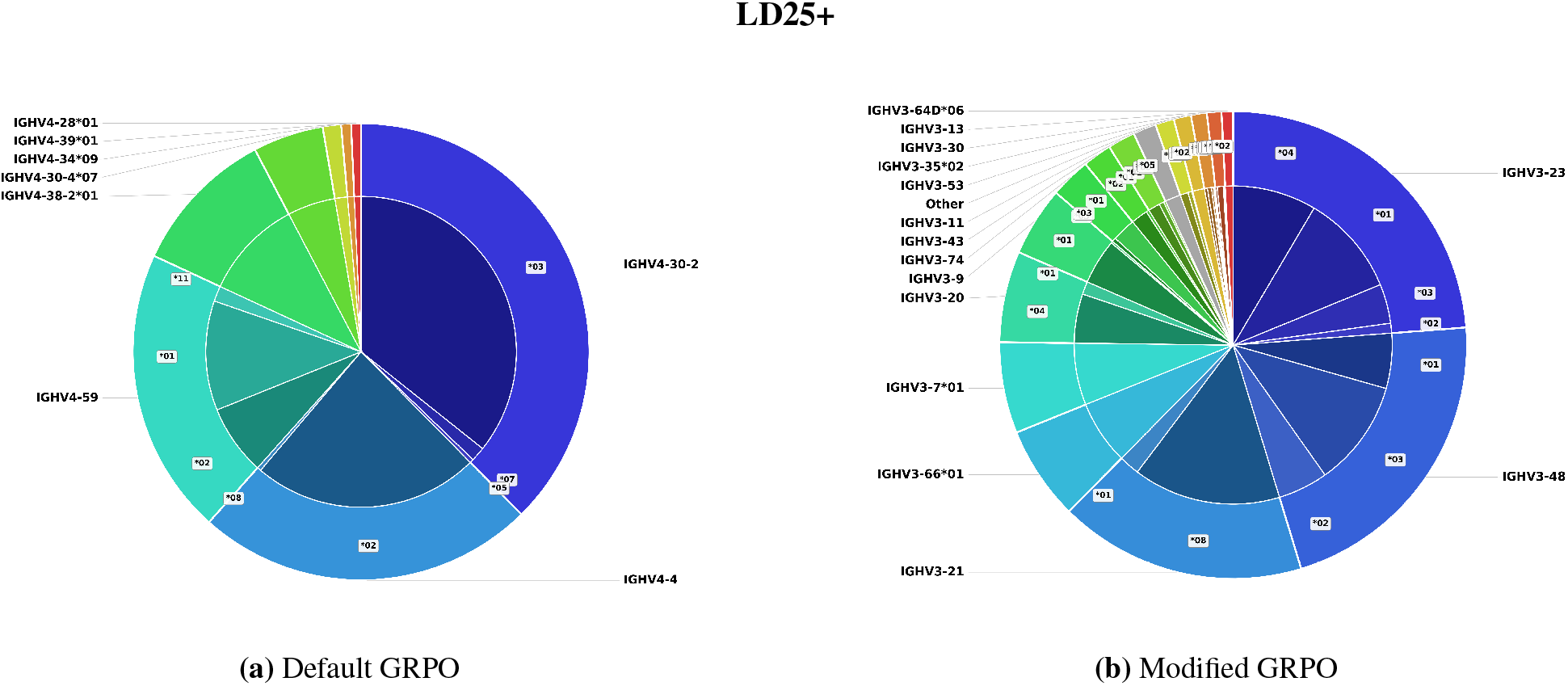
Distribution of unique germline V calls at LD25 threshold.

**Figure E.5:**
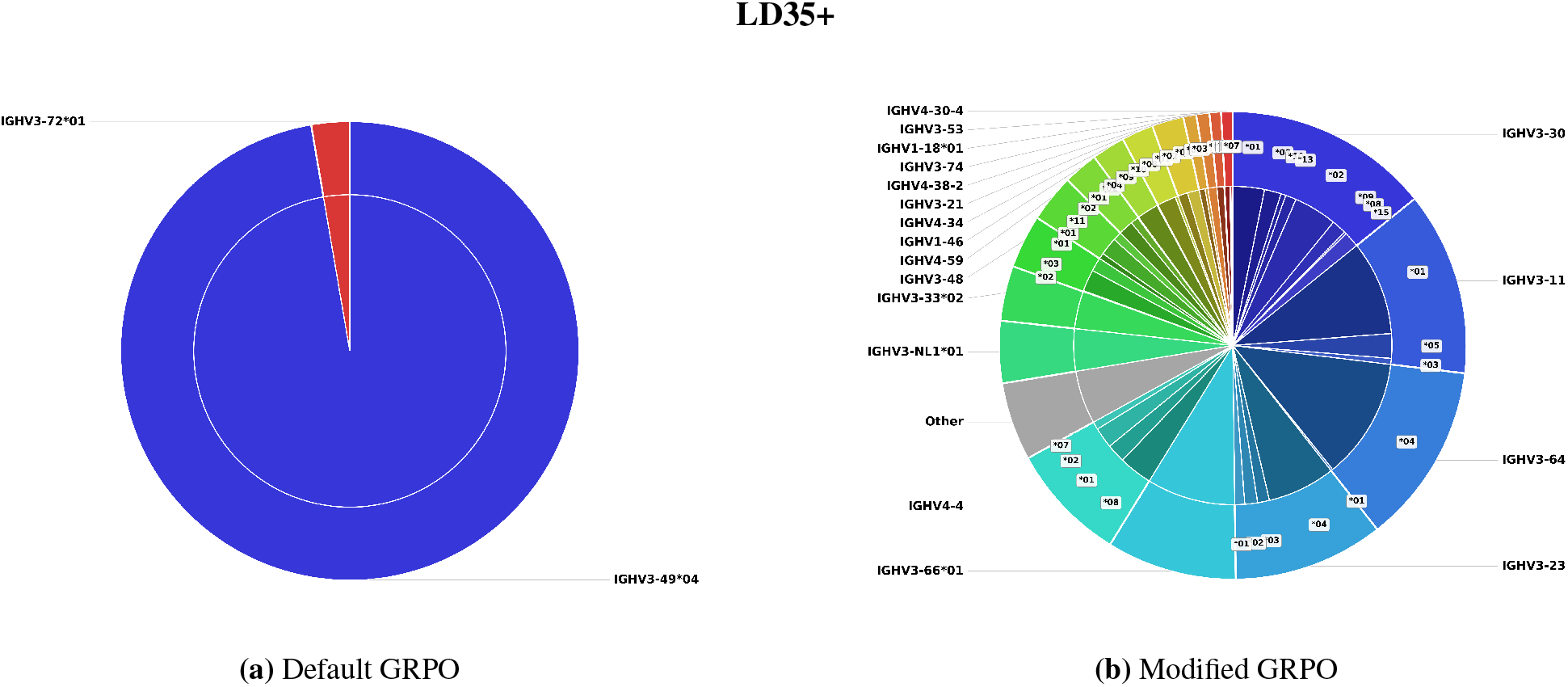
Distribution of unique germline V calls at LD35 threshold.

##### E.2.2 Unique J calls

**Figure E.6:**
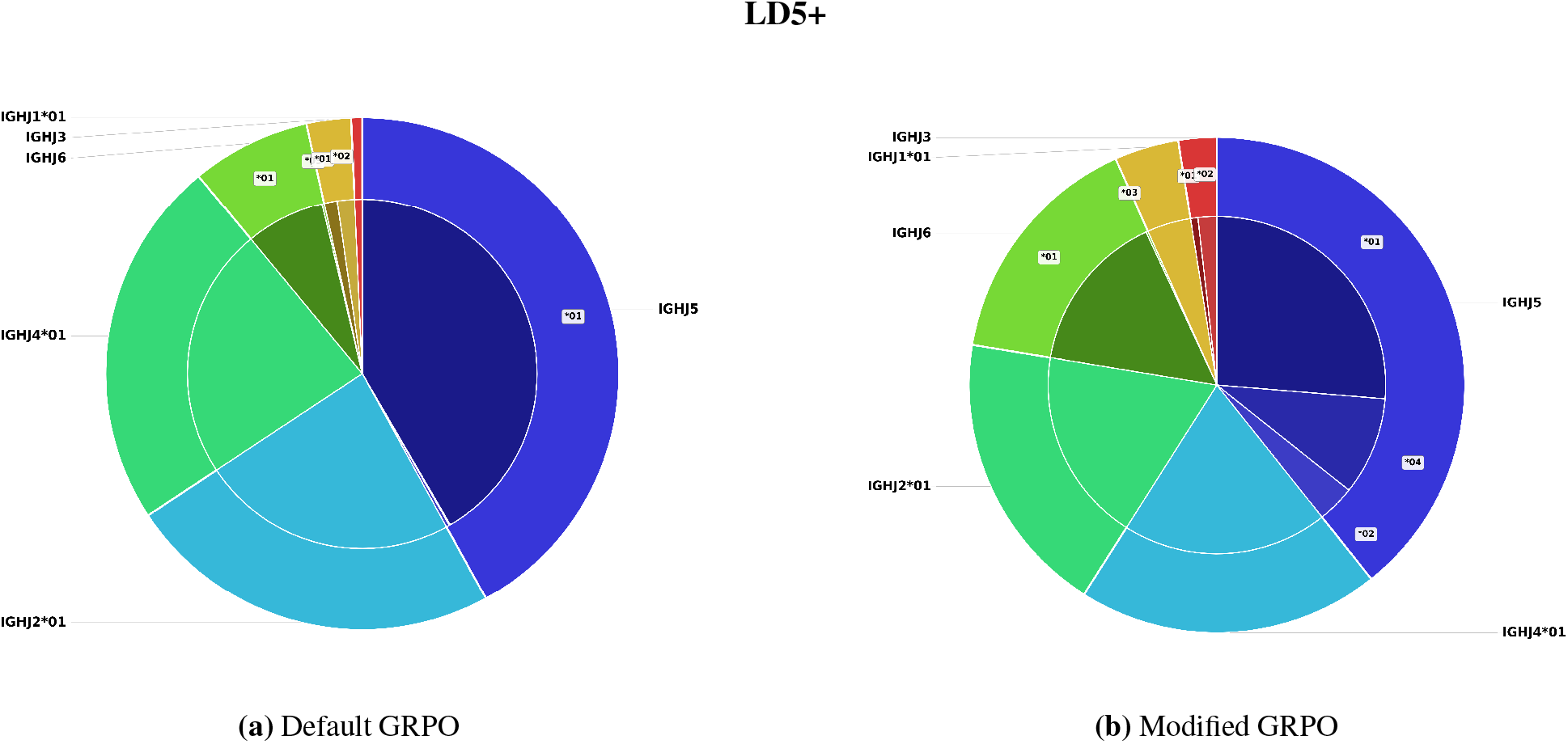
Distribution of unique germline J calls at LD5 threshold.

**Figure E.7:**
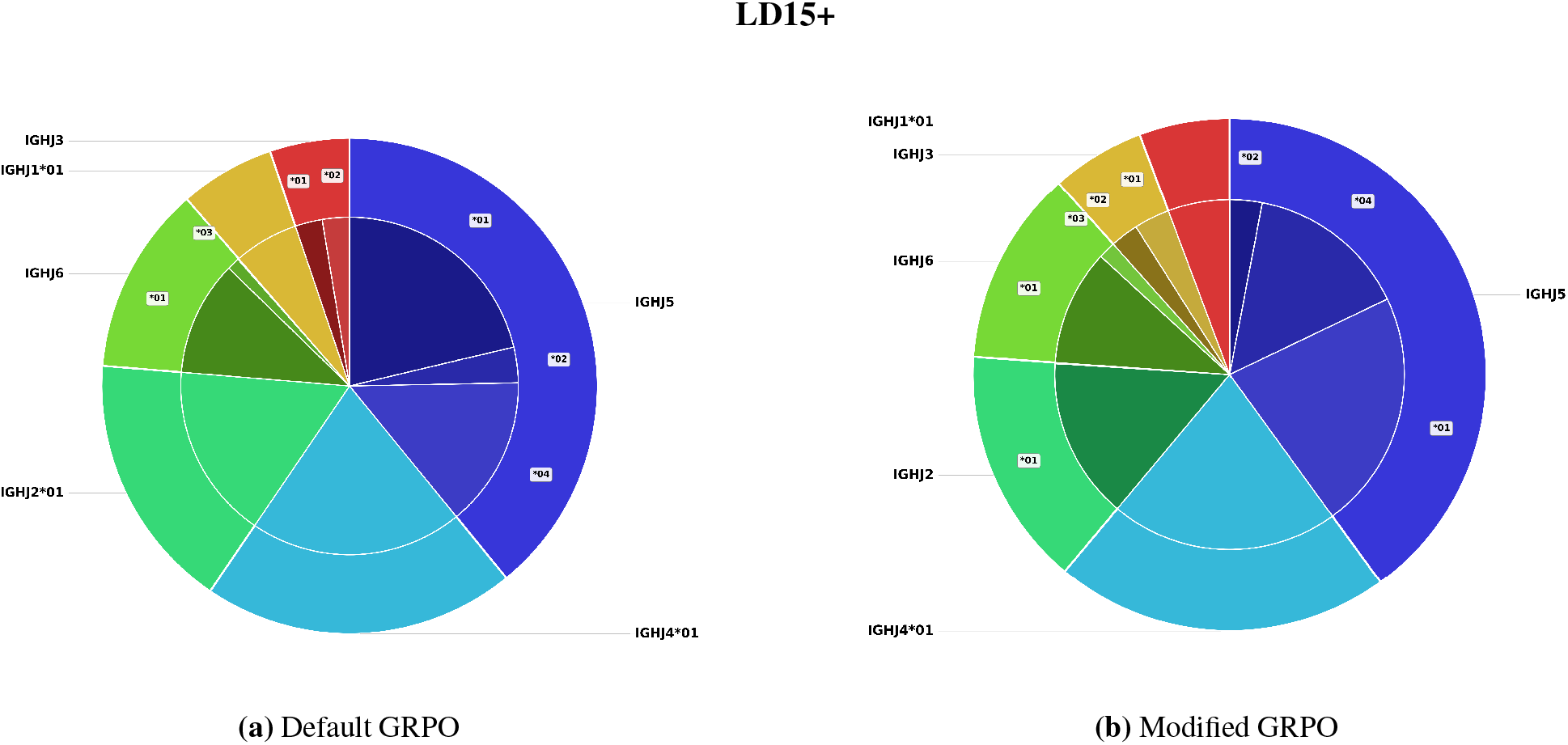
Distribution of unique germline J calls at LD15 threshold.

**Figure E.8:**
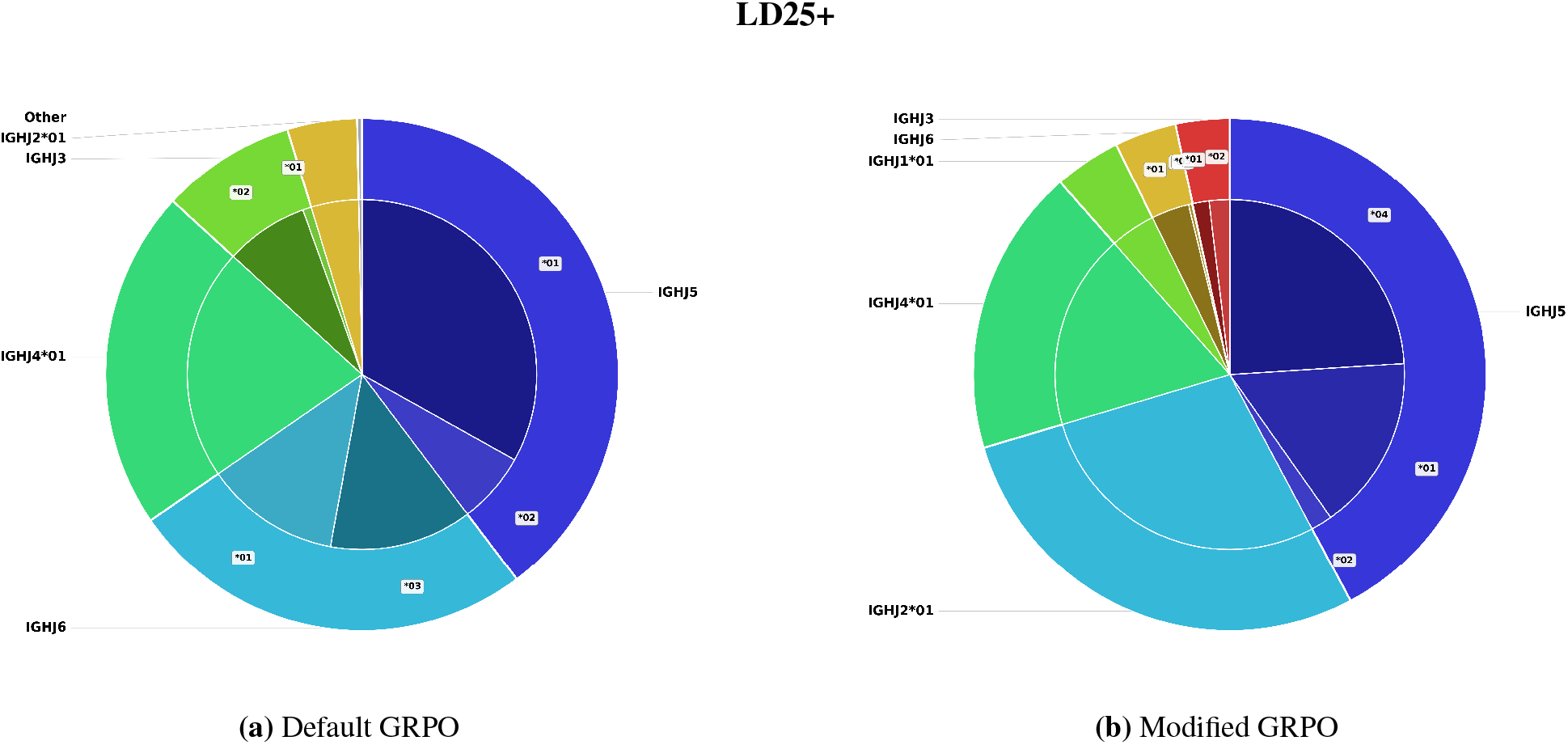
Distribution of unique germline J calls at LD25 threshold.

**Figure E.9:**
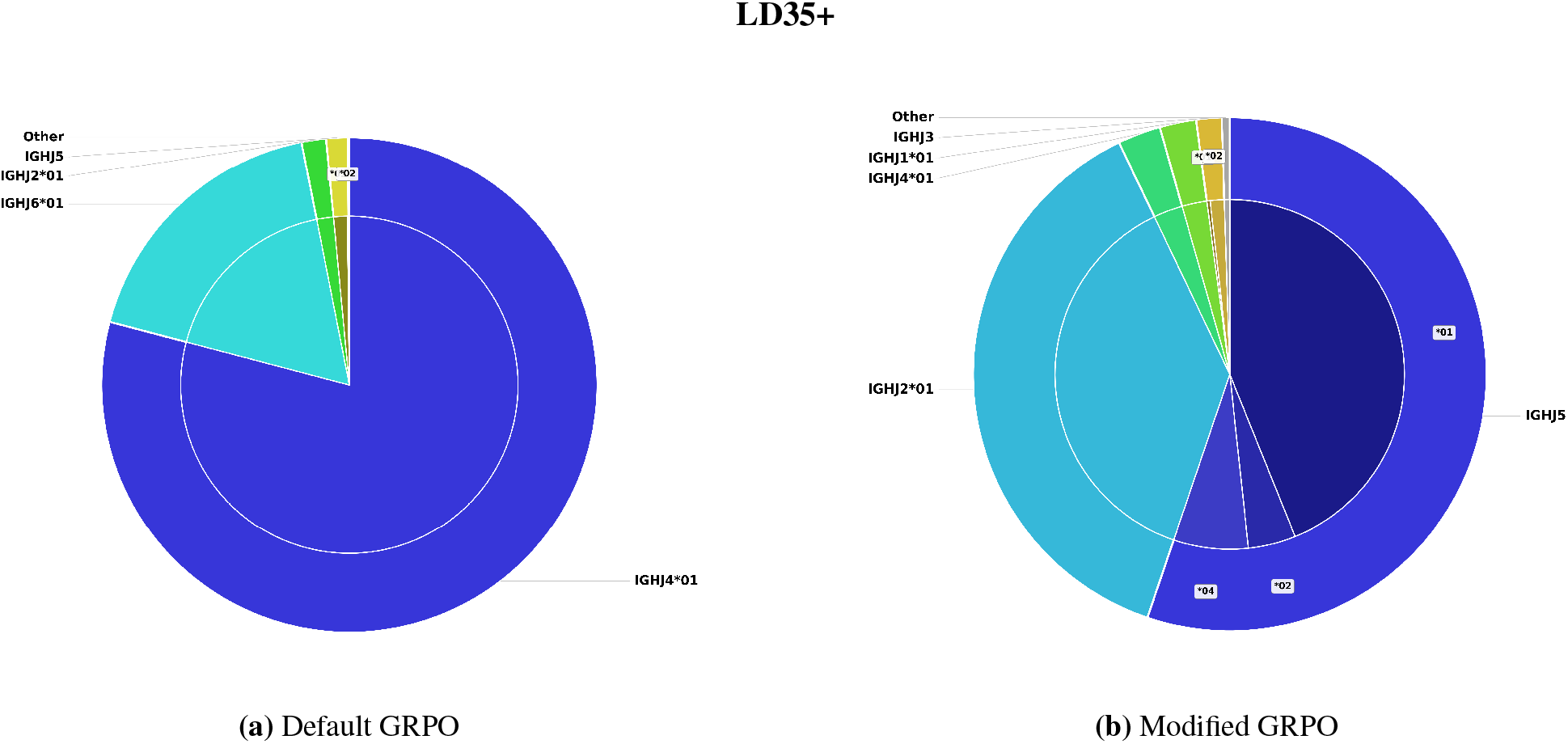
Distribution of unique germline J calls at LD35 threshold.

#### E.3 Bounded LD_Thresh_ normal and RGT reward training unique germline calls from sequence generations after RL training

##### E.3.1 Unique V calls

**Figure E.10:**
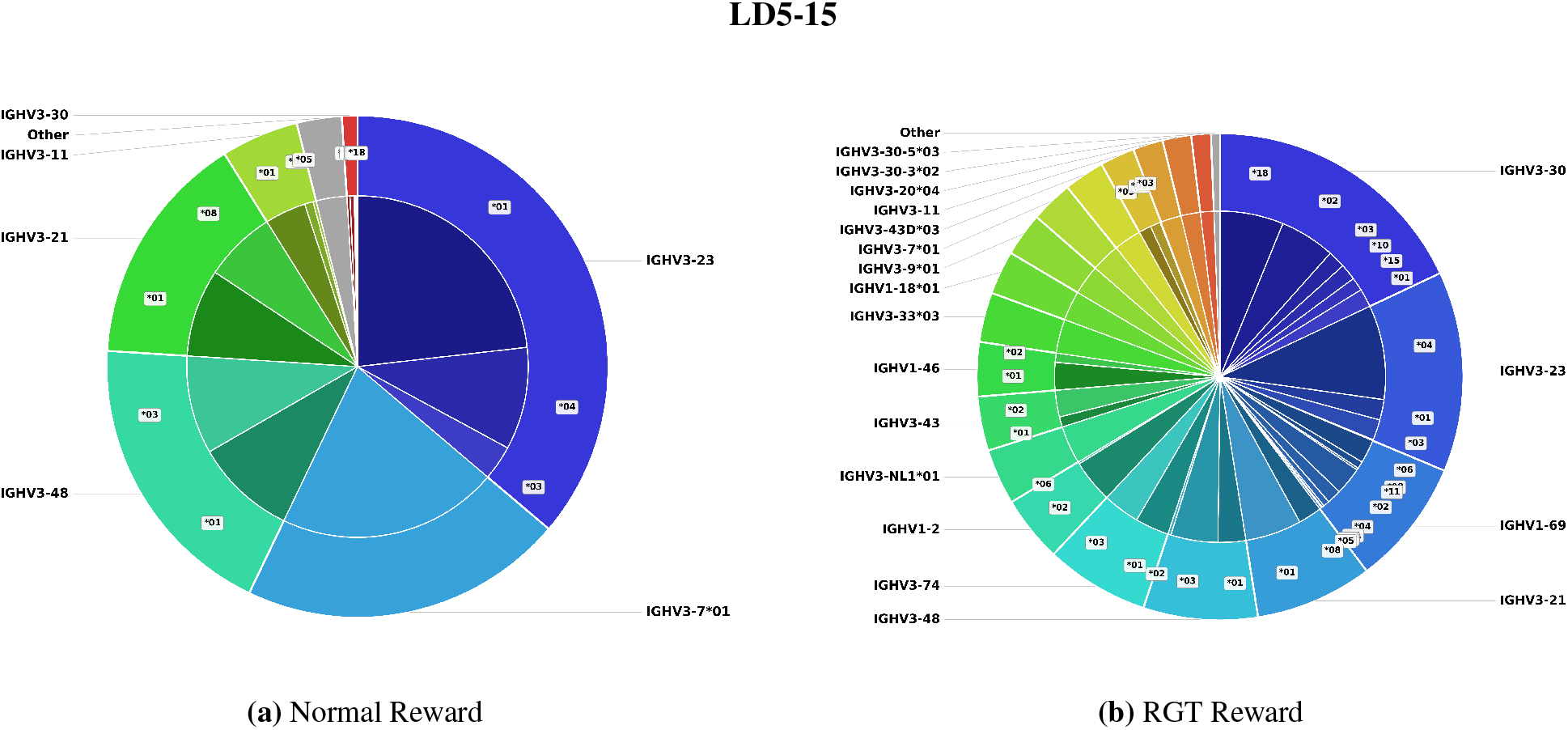
Distribution of unique germline V calls at bounded LD5–15 threshold under normal and RGT rewards.

**Figure E.11:**
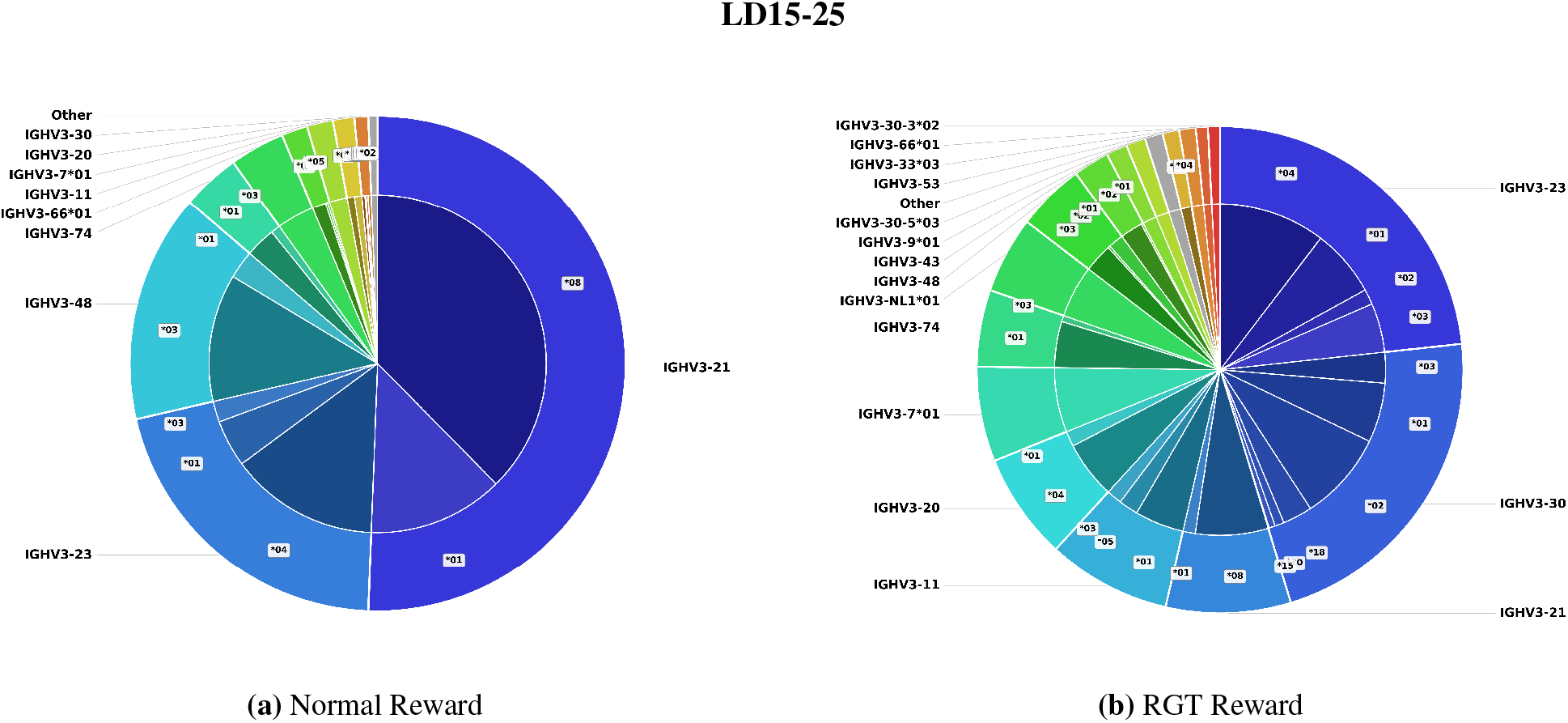
Distribution of unique germline V calls at bounded LD15–25 threshold under normal and RGT rewards.

**Figure E.12:**
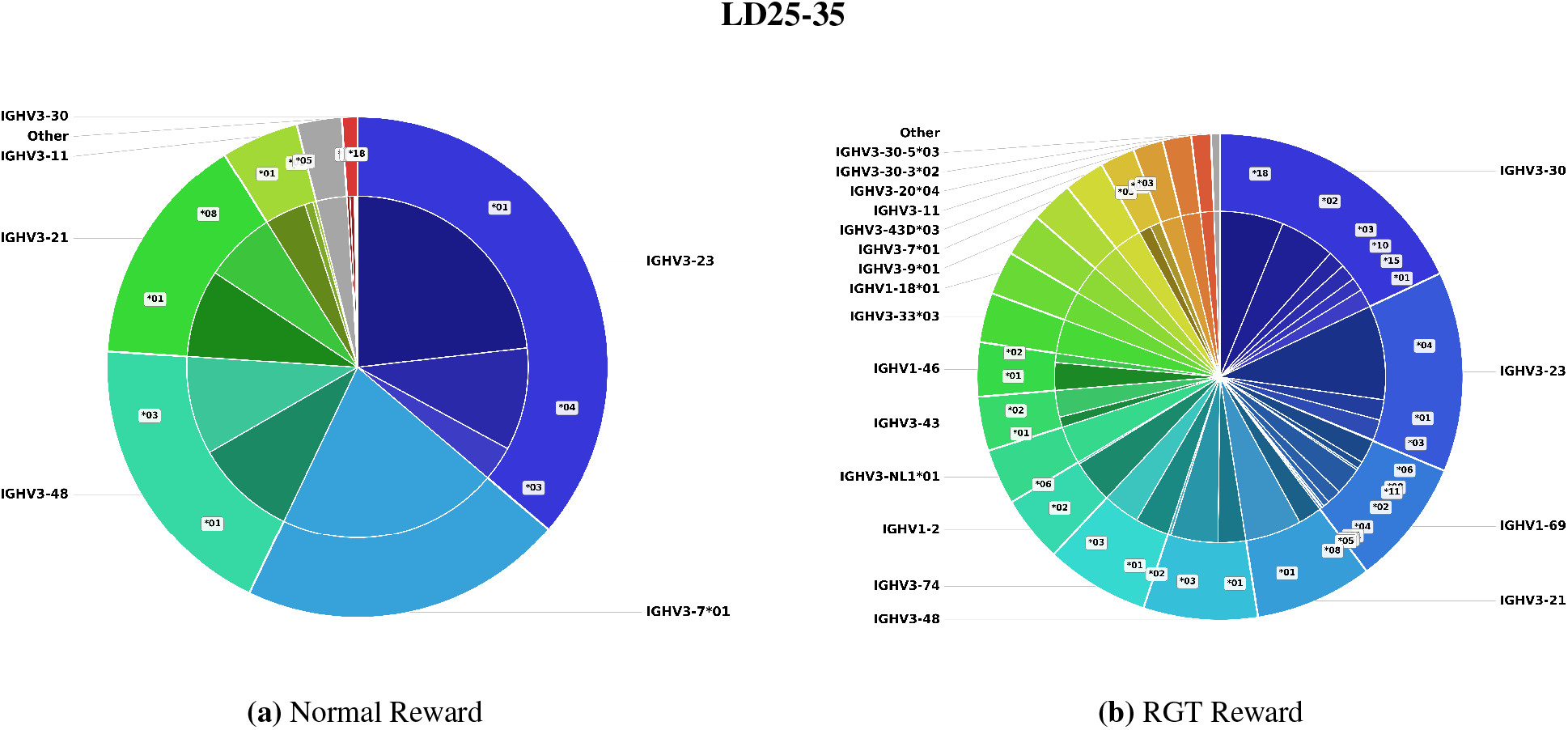
Distribution of unique germline V calls at bounded LD25–35 threshold under normal and RGT rewards.

##### E.3.2 Unique J calls

**Figure E.13:**
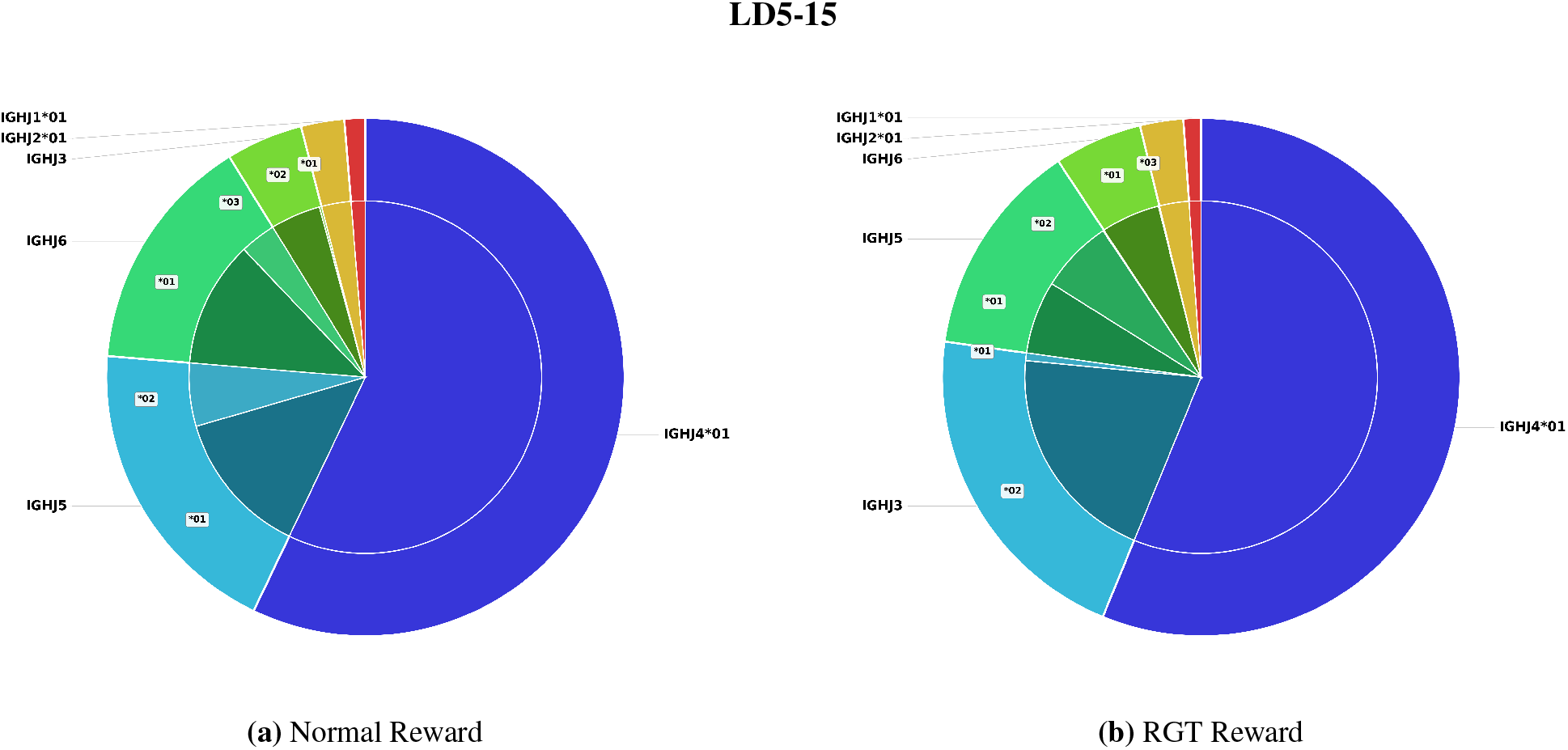
Distribution of unique germline J calls at bounded LD5–15 threshold under normal and RGT rewards.

**Figure E.14:**
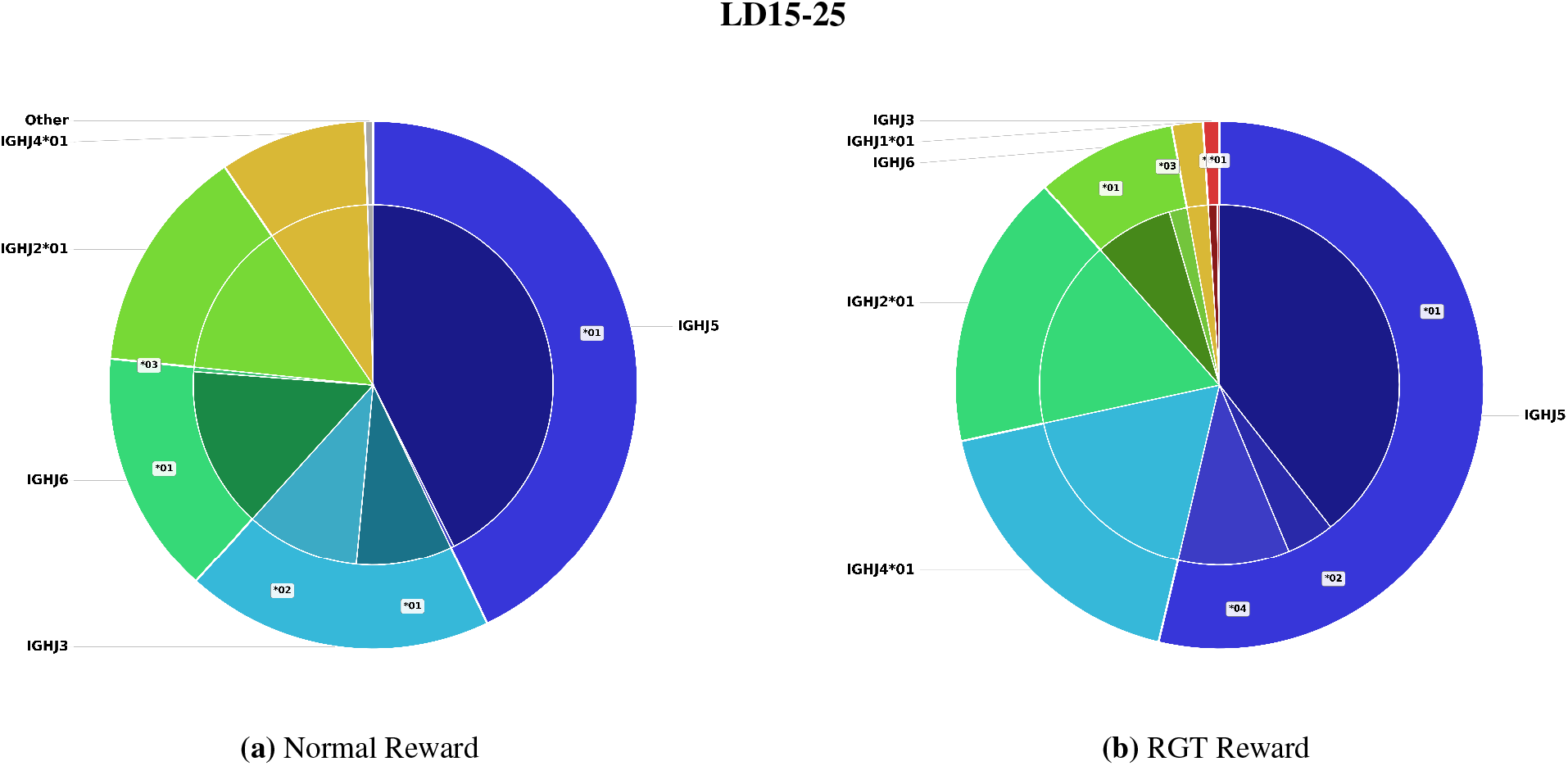
Distribution of unique germline J calls at bounded LD15–25 threshold under normal and RGT rewards.

**Figure E.15:**
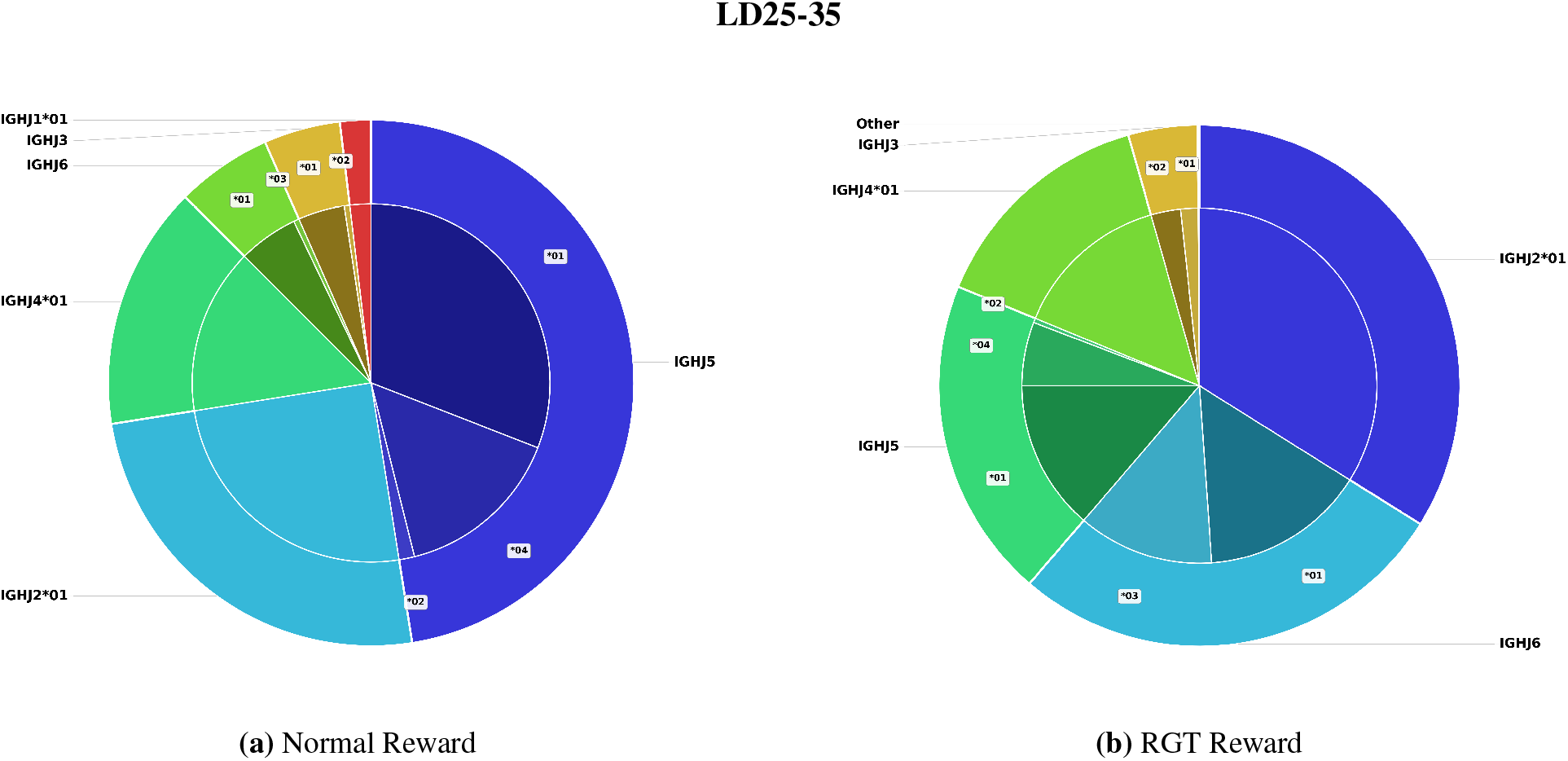
Distribution of unique germline J calls at bounded LD25–35 threshold under normal and RGT rewards.

### F Customization and control with bound LD objectives with GermRL

RL training with the single LD threshold objective successfully generated high LD antibodies that maintain strong sequence folding confidence to alleviate the germline bias. However, without an upper LD bound, the policy is unrestricted in the LD distance it generates. In antibody engineering efforts, it is practical to target mutations within a certain range in order to promote diversity without straying too far from germline. Therefore, we ask if it is possible to constrain the LD of antibody generations during training to control the extent of mutations within a generated sequence.

#### F.1 Addition of upper bound to LD reward constrains distance from germline

To bound antibody LD generations, we add an upper threshold LD bound where the reward decays as the LD of the generated sequence exceeds the upper bound (**Appx. A.2**). Given the additional complexity of this objective, we observe the policy sampling from a small pool of germline V calls at certain LD bound objectives, shown in **Fig. F.2**. To prevent this behavior, we also introduce another reward component alongside the bounded LD reward, named “Reward Germline Trajectory” (RGT), which penalizes the policy from continuously generating from overexplored germline calls (**Appx. A.2**).

To investigate the success of adding the LD constraint to the reward function with and without RGT, we apply GermRL to train 3 versions of ProGen2-OAS with different LD threshold bounds: LD5-15, LD15-25, LD25-35, using modified GRPO. After training, we generate 1,000 antibody sequences from each policy and evaluate their pass@1 for the success condition of falling within the LD threshold bounds and the condition of falling within the LD threshold bounds and surpassing the pLDDT threshold of 0.7. The pass@1 values of the RL trained policies are shown in **Table F.1** while the scatter plots of LD and ESMFold pLDDT for each generation are shown in **Fig. F.1**.

**Table F.1:** Probability of one-shot success (pass@1) between Normal Reward and RGT Reward across LD bins under LD-only (top 3 rows) and LD with foldability (bottom 3 rows) success conditions for the bounded LD objective.

| Condition | LD | GermRL Bound Normal Reward |  |  | GermRL Bound RGT Reward |  |  |
| --- | --- | --- | --- | --- | --- | --- | --- |
|  |  | LD5-15 | LD15-25 | LD25-35 | LD5-15 | LD15-25 | LD25-35 |
| $LD_{\text{thresh}_{\text{high}}} \geq LD \geq LD_{\text{thresh}_{\text{low}}}$ | 5-15 | **0.960** | 0.062 | 0.000 | **0.960** | 0.049 | 0.000 |
|  | 15-25 | 0.070 | **0.961** | 0.059 | 0.039 | 0.913 | 0.183 |
|  | 25-35 | 0.000 | 0.020 | **0.927** | 0.001 | 0.106 | 0.885 |
| $LD_{\text{thresh}_{\text{high}}} \geq LD \geq LD_{\text{thresh}_{\text{low}}}$<br>and pLDDT > 0.7 | 5-15 | **0.960** | 0.062 | 0.000 | 0.959 | 0.049 | 0.000 |
|  | 15-25 | 0.070 | **0.961** | 0.059 | 0.039 | 0.912 | 0.183 |
|  | 25-35 | 0.000 | 0.020 | **0.921** | 0.001 | 0.105 | 0.876 |

**Figure F.1:**
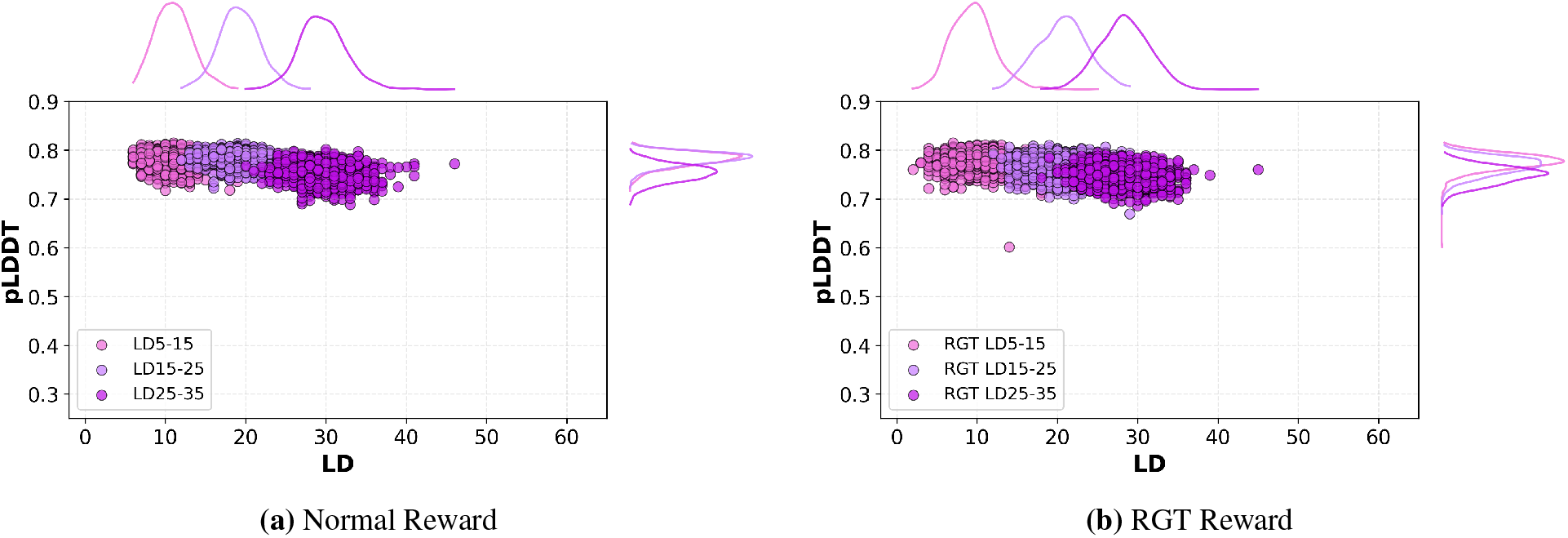
LD and pLDDT distribution from ProGen2-RL policies under the bounded LD objective with the normal reward and RGT reward.

From **Table F.1**, policies trained on a specific bounded objective learn to generate within the bounded region. Although the normal reward achieves higher or equal pass@1 for each bounded LD objective compared to the RGT reward, training with the RGT reward maintains a steady generation of unique germline calls within the batch of trajectories for each training step in an epoch (**Fig. F.2**).

Through the sequence similarity and cumulative density plot of top-*k* germline calls from the set of 1,000 antibodies generated post training, shown in **Fig. F.3**, and pieplots of the germline call distribution, shown in **Appx. E**, we demonstrate the application of the RGT reward encouraging a greater equilibrium in the distribution of generated germline calls, lowering the sequence similarity of the generations. However, the distribution of explored germline calls after RL training with the bounded LD objective exhibits reduced diversity compared to base ProGen2-OAS and under RL training with the single LD objective (**Appx. E**), establishing room for future improvement of GermRL. For our analysis in Section 6, we use the GermRL bounded objective with the RGT reward.

**Figure F.2:**
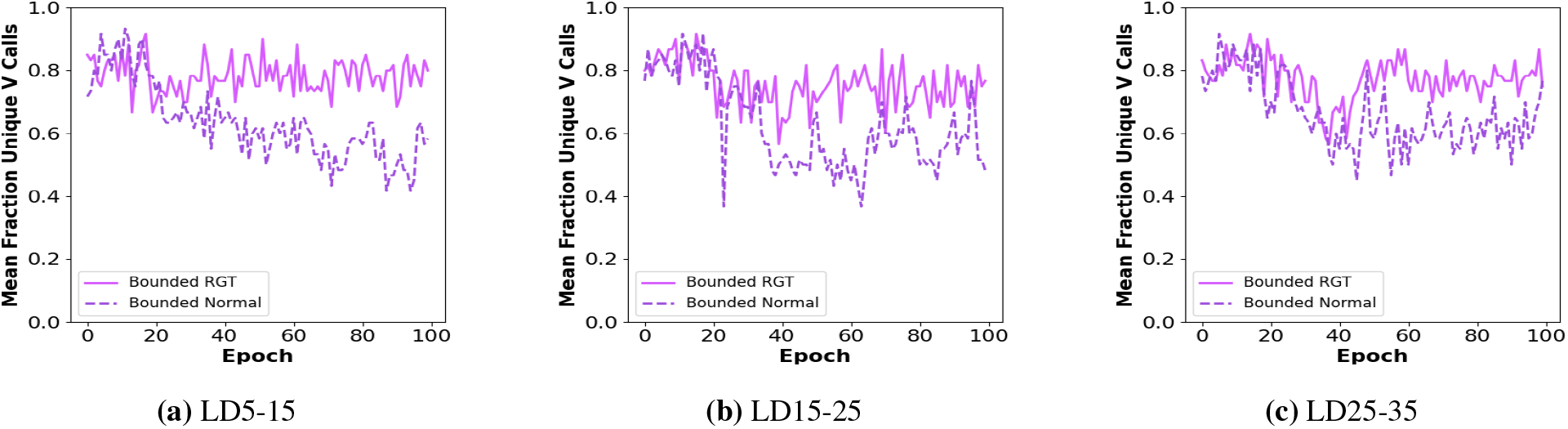
Mean fraction of generated antibodies with unique germline V calls within the trajectory batch for a training step across all epochs of RL training between normal and RGT reward for the bounded LD objective.

**Figure F.3:**
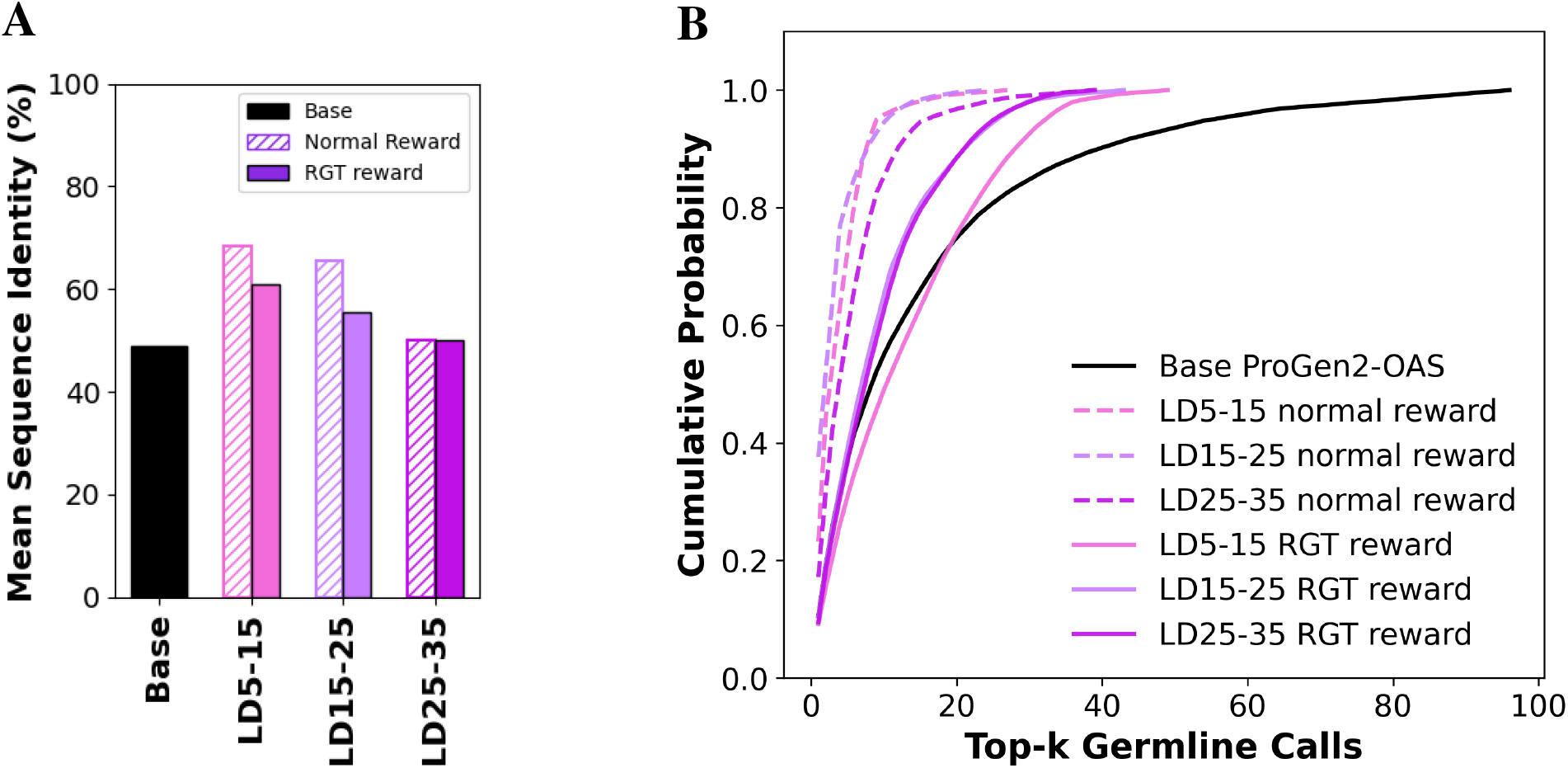
**(a)** Mean sequence similarity between each pair of antibodies from all base ProGen2-OAS generations and policy generations after RL training with modified GRPO under the bounded LD objective for the normal and RGT reward. **(b)** Cumulative distribution of top-*k* germline V calls of generated sequences from base ProGen2-OAS after RL training between normal and RGT reward for the bounded LD objective.

### G UMAP J call projection

**Figure G.1:**
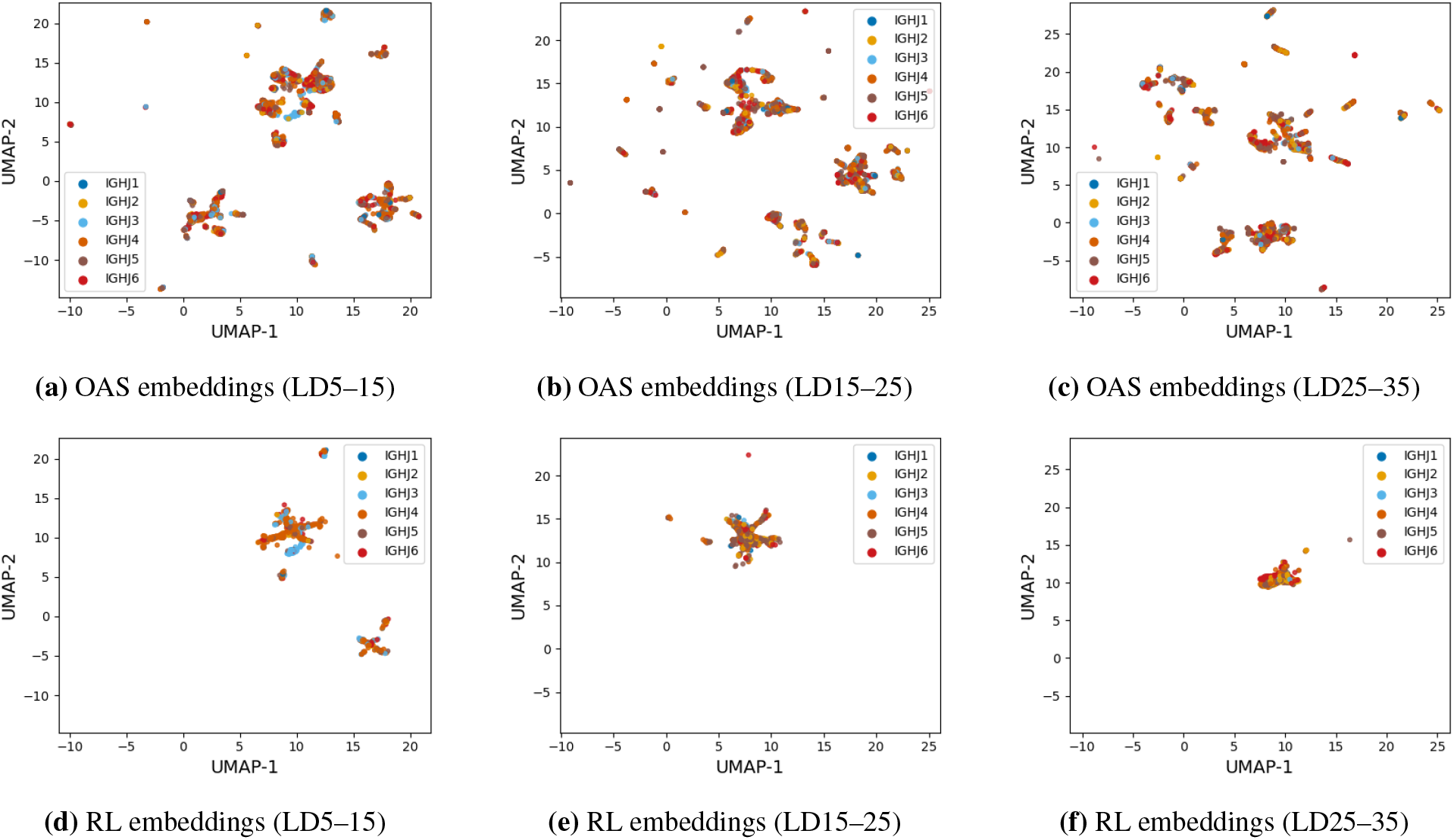
UMAP embeddings colored by J gene family across different Levenshtein distance (LD) ranges. (**a**–**c**) OAS embeddings for LD5–15, LD15–25, and LD25–35. (**d**–**f**) RL-generated embeddings for LD5–15, LD15–25, and LD25–35.

### H Base ProGen2-OAS and OAS biophysical property distribution analysis

To quantify the extent of antibody property divergence seen in RL-generated antibodies attributed to ProGen2-OAS pre-training, we repeat all analysis presented in Section 6 with base ProGen2-OAS. To sample high LD antibodies from the base AbLM, we use a temperature of 1.3 and retain only generated antibodies that surpass pLDDT > 0.7, resulting in 607 sequences between 5-25 LD. The specific distribution of LD sequences between the bound are shown in **Fig. H.2b** where the majority of generations fall under 12 LD. We sample 5,000 OAS sequences between 5-25 LD to match this specific LD distribution of base antibodies.

#### H.1 Base ProGen2-OAS and OAS regional mutation distribution analysis

**Figure H.1:**
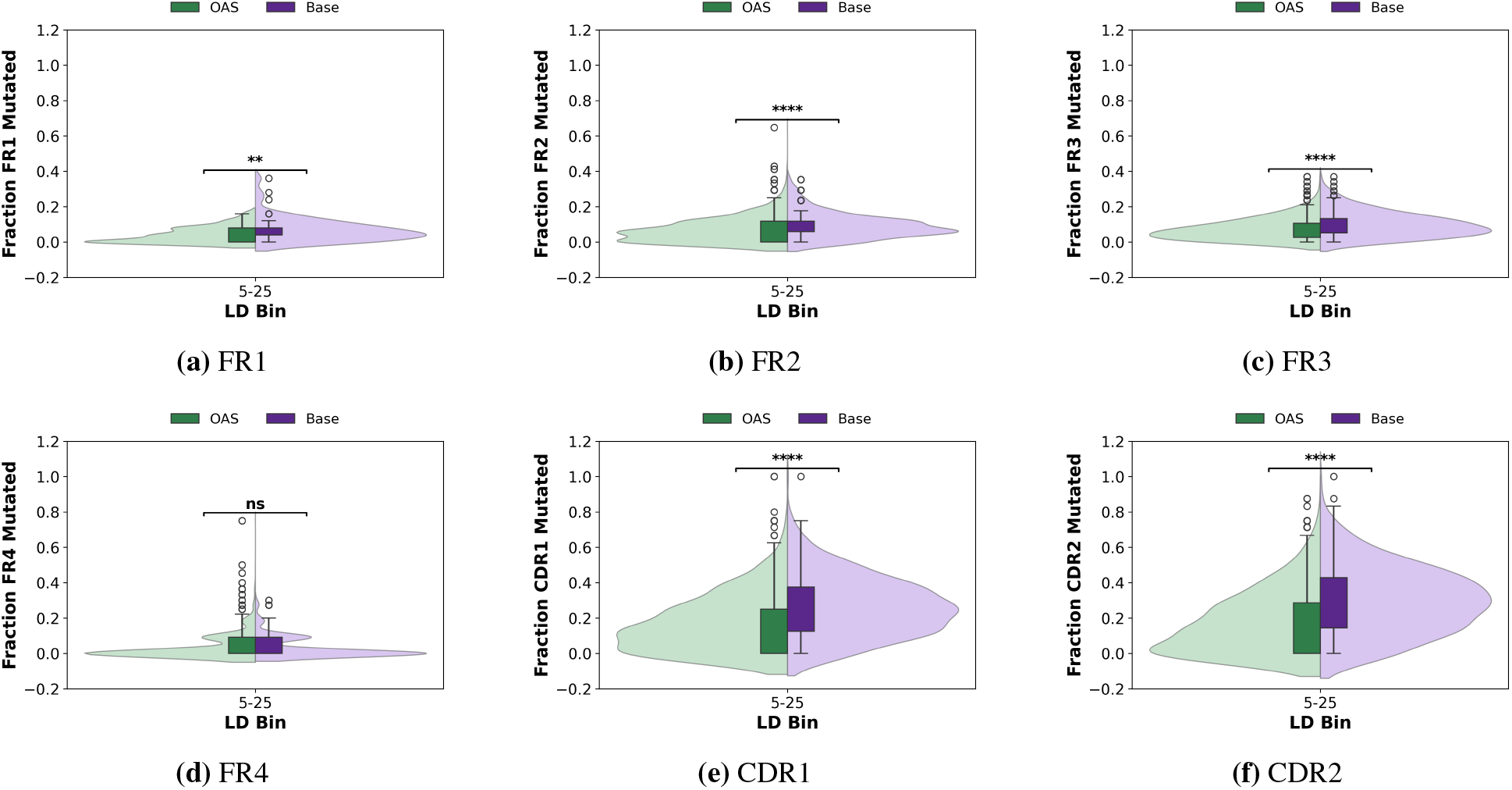
Fraction of each antibody region that is mutated for base ProGen2-OAS generated and OAS sequence

**Figure H.2:**
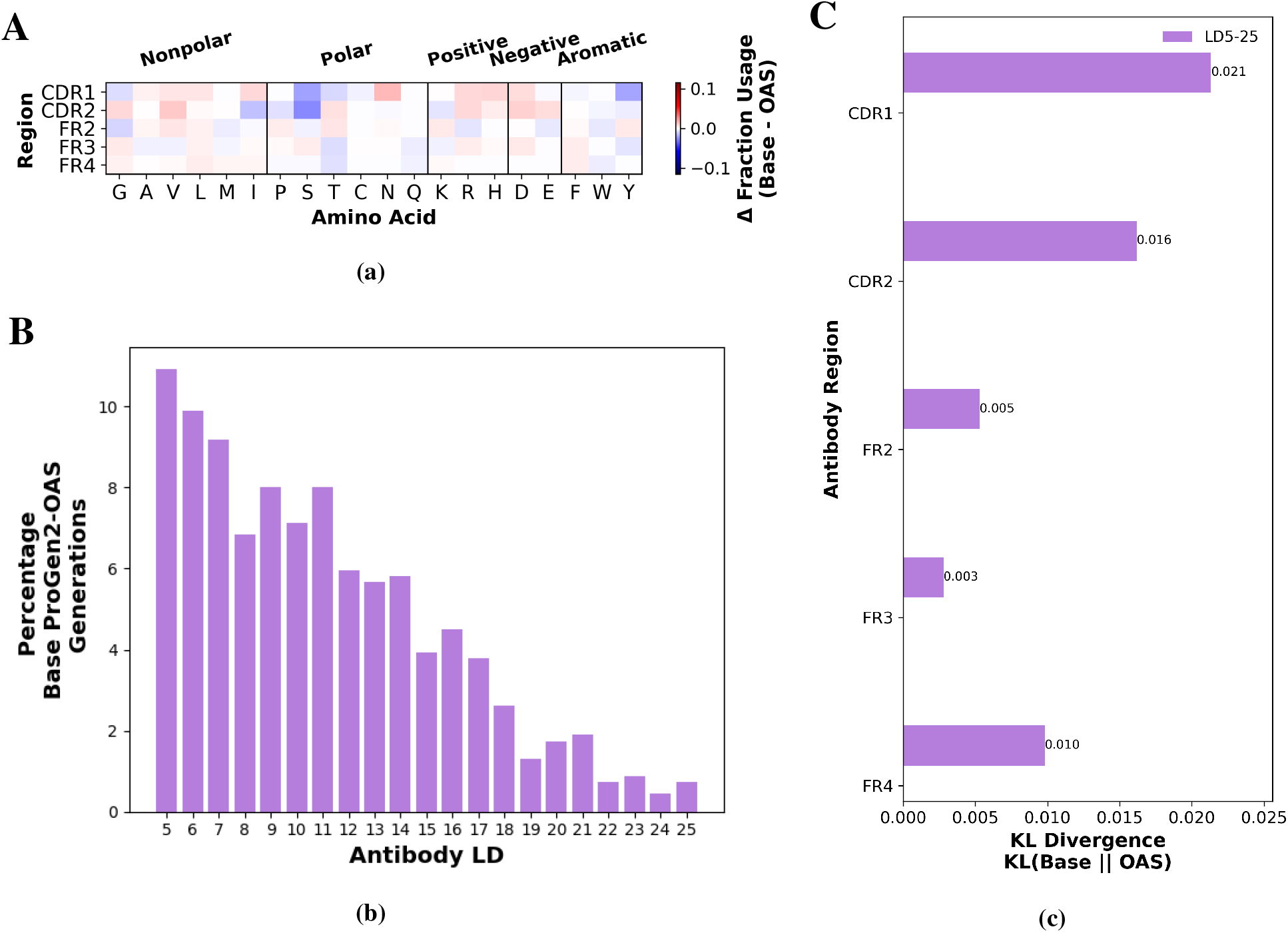
(**a**) Breakdown of difference in regional amino acid fractional composition between base ProGen2-OAS and OAS sequences across LD 5–25. (**c**) LD distribution of 607 Base ProGen2-OAS generated antibodies for OAS comparison analysis. (**c**) KL divergence between base ProGen2-OAS and OAS amino acid distributions for increasing LD ranges.

#### H.2 Base ProGen2-OAS and OAS BLOSUM score distribution analysis

**Figure H.3:**
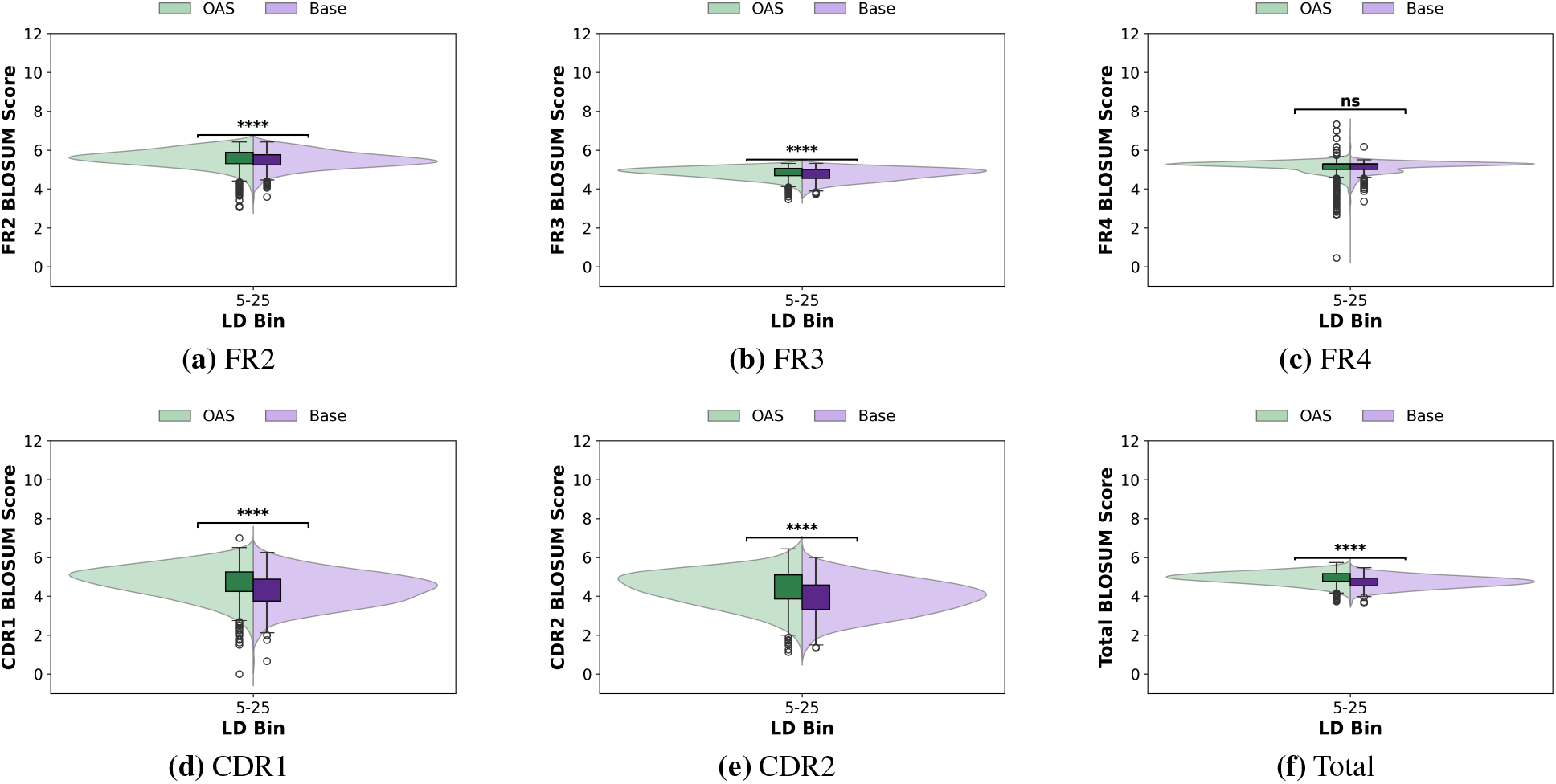
Mean BLOSUM62 scores from base ProGen2-OAS and OAS sequences across LD bins and antibody regions

#### H.3 Base ProGen2-OAS and OAS developability analysis

**Figure H.4:**
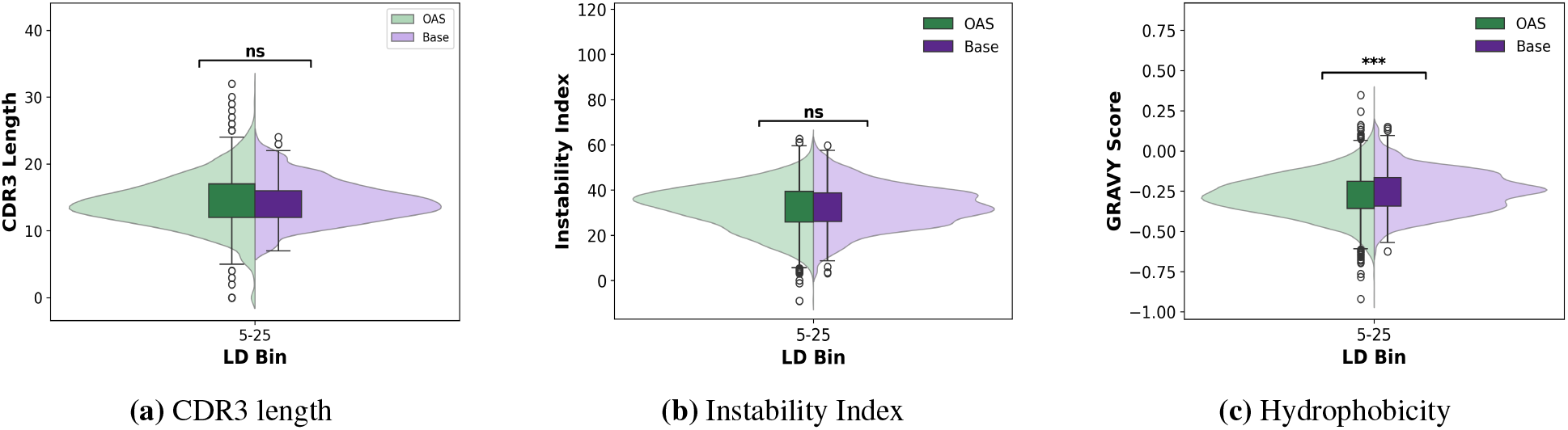
**(a)** CDR3 length, **(b)** Instability Index and **(c)** Hydrophobicity GRAVY score between base ProGen2-OAS generated and OAS sequence.

**Figure H.5:**
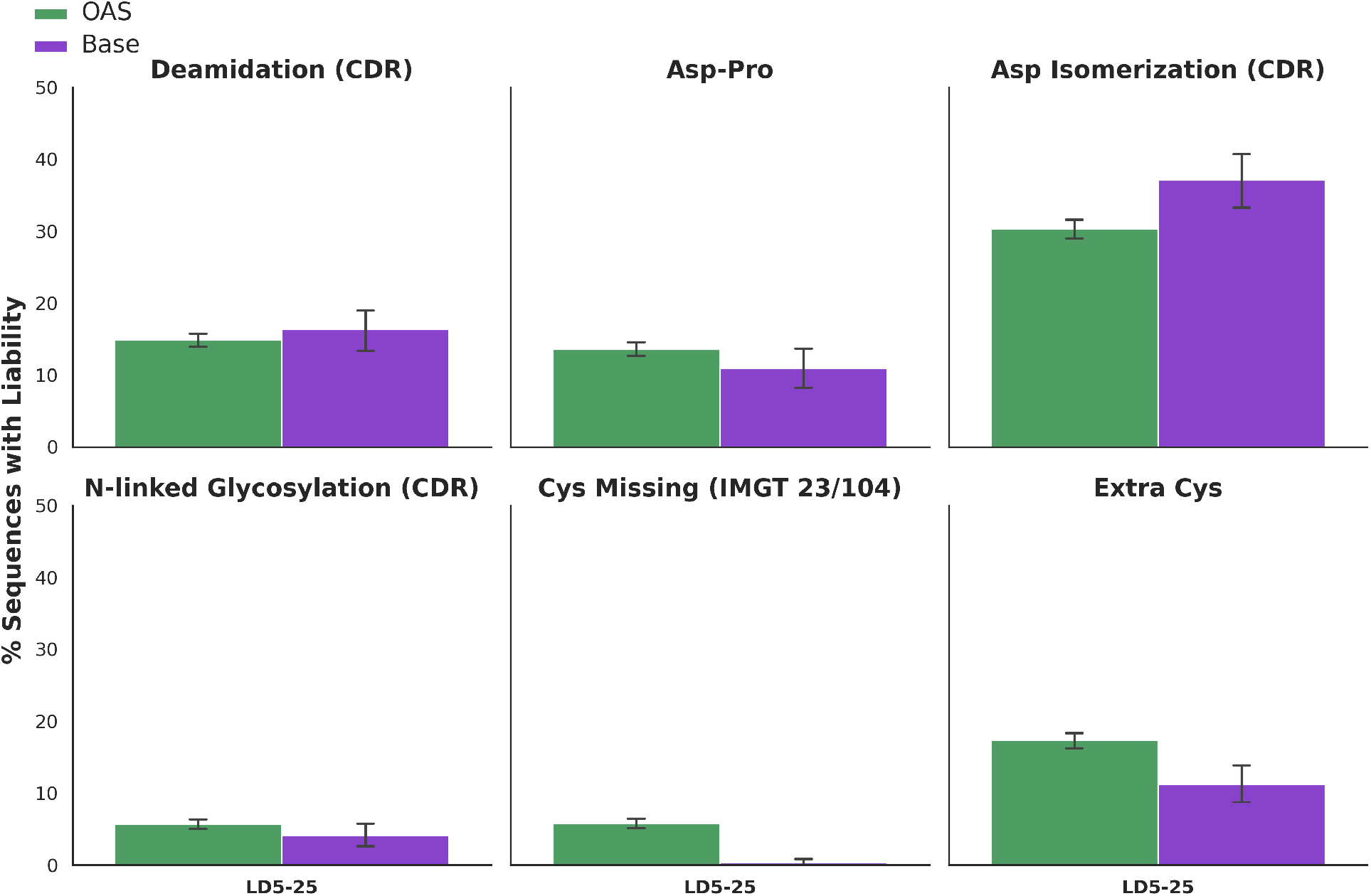
Percent of sequences containing liable motifs between base ProGen2-OAS generated and natural OAS sequences. Asp isomerization, deamidation and n-linked glycosylation liabilities are searched in each of the 3 CDR regions.

